# Breeding value theory with genetically differentiated mate pools and imprinting

**DOI:** 10.64898/2026.11.23.767970

**Authors:** David López-Carbonell, Luis Varona, R. Chris Gaynor, Gregor Gorjanc

**Affiliations:** Facultad de Veterinaria, Instituto Agroalimentario de Aragón (IA2), Universidad de Zaragoza, 50013, Zaragoza, Spain; The Roslin Institute and Royal (Dick) School of Veterinary Studies, University of Edinburgh, Easter Bush, Edinburgh, EH25 9RG, Midlothian, United Kingdom; Bayer Crop Science, 700 Chesterfield Pkwy W., Chesterfield, MO 63017, Missouri, USA

**Keywords:** Breeding value, Additive genetic value, General combining ability, Imprinting, Genetically differentiated mate pools

## Abstract

Breeding value is commonly identified with additive genetic value. These measures of genetic merit are equivalent when the reference population and mate pools share the same allele frequency, and there are no parent-of-origin effects. In breeding programs, however, female and male mate pools can differ in allele frequency, and imprinting can affect the expected genetic value of their progeny. We review the standard one-locus model with additive and dominance effects under genetically undifferentiated and differentiated mate pools, and repeat the analysis with imprinting effects. We distinguish additive genetic value, defined by the within-generation regression of genetic value on allele dosage, from breeding value, defined by the regression of expected progeny genetic value on parental allele dosage. Under this definition, breeding value is twice the slope of between-generation regression multiplied by centered parental allele dosage. The Price equation connects these breeding-value expressions to the change in mean genetic value under selection. Under Hardy–Weinberg genotype frequencies, with undifferentiated mate pools and no imprinting, the two additive genetic and breeding values coincide. They can differ with dominance and differentiated mate pools, or with imprinting. We show that in these models, general combining ability equals one-half of breeding value when both are defined for the same mate population and relative to the same mean progeny value. We illustrate these relationships with a one-locus example and use a multi-locus simulation to examine allele frequency differences under antagonistic and synergistic selection. An analysis of Rubia Gallega cattle data shows that allele frequency can differ between female and male reproductive groups, highlighting the relevance of the nuanced definition of breeding value in breeding programs. This work clarifies how the choice of reference population, mate pool, and parental role affects the interpretation of genetic merit in selective breeding.

## Introduction

**S**elective breeding is based on identifying and using the genetically best individuals as parents of the next generation. Historically, selection decisions were based directly on pheno-typic values, whereas modern selective breeding uses statistical estimates of genetic merit obtained from phenotypic and pedigree data (Henderson 1984; Mrode and Pocrnic 2023), and, increasingly, genomic data (Meuwissen et al. 2001). By better separating the genetic and environmental components of observed phenotypic values, selection and mating based on estimated genetic merit increase the rate of genetic improvement for complex traits relative to phenotype-based selection (Lush 1943; Falconer 1981). Over time and different contexts, several related quantities have been defined to quantify the genetic merit of selection candidate as a parent. Most common are additive genetic value (Fisher 1919, 1930, 1958), breeding value (Lush 1943), and general combining ability (GCA) (Sprague and Tatum 1942; Griffing 1956b,a).

Although these quantities are well established, their interpretation depends on reference populations and mating assumptions that are often left implicit. These assumptions become particularly important when reproductive groups differ in allele frequency or when allele effects depend on parental origin. In the breeding context, female and male candidates for reproduction can be considered as two different mating pools. We distinguish two settings: genetically undifferentiated mate pools in which the female and male pools share the reference population’s allele frequency, and genetically differentiated mate pools, in which the female and male pools differ in allele frequency. Hereafter, we refer to these respectively as undifferentiated and differentiated mate pools.

Our aim is to clarify the relationships among additive genetic value, breeding value, and GCA under undifferentiated and differentiated mate pools, without and with imprinting. We review their definitions and assumptions, distinguishing the withingeneration regression defining additive genetic value from the between-generation regression defining breeding value. Using one-locus models, we examine when additive genetic value and breeding value coincide or differ, and how mate-pool-specific breeding value relates to GCA. We then examine differences in female and male allele frequencies through simulation and data from Rubia Gallega beef cattle to illustrate the relevance of these 2 Breeding value theory under different conditions distinctions for selective breeding.

### Review

Fisher’s statistical decomposition of phenotypic values into genetic values and environmental deviations provides the framework to define these quantities (Fisher 1919, 1930, 1958). These genetic values are due to genotype differences between individuals and additive (allele) effects as well as non-additive (interaction) effects such as dominance (interaction) effects between alleles at one locus, epistatic (interaction) effects between genotypes at different loci, and imprinting (interaction) effects of alleles due to their parental origin (Falconer 1981; Lynch and Walsh 1998; Walsh and Lynch 2018). He further statistically decomposed an individual’s genetic value into an additive genetic value and a non-additive genetic deviation by regressing genetic value on the number of alternative alleles (allele dosage). Points on this regression line, expressed as deviations from the population mean, represent additive genetic values, while deviations from it represent non-additive genetic deviations. Using this decomposition, he derived expected correlations between different pairs of relatives.

The slope of this regression (*α*) is therefore the essential quan-tity. Although additive genetic values are linear in allele dosage, they can reflect part of the underlying non-additive effects because these effects can influence *α* (Falconer 1981; Lynch and Walsh 1998; Hill et al. 2008; Hivert et al. 2021). Since this regression is defined with respect to the genotype distribution in a reference population, changes in genotype frequencies can change *α* and hence both the corresponding additive genetic values and the portion of non-additive effects that they capture, making additive genetic values population-specific (Falconer 1981; Lynch and Walsh 1998; Hivert et al. 2021).

Population specificity becomes especially important when genetic merit is interpreted across generations and mate pools, as in the concept of breeding value. This term arose in practical animal breeding (e.g. Lippincott 1923; Davidson 1925) and was formalized by Lush (1935, 1943) as an individual’s genetic merit as a parent, operationally expressed as twice the difference between the individual’s expected progeny mean and the corresponding population progeny mean. Related measures on the one-half-breeding-value scale include transmitting ability and expected progeny difference (Yapp 1925; Beef Improvement Federation 1972; Falconer 1981). Lush connected this progeny-based definition to the regression theory of Fisher (1919) and Wright (1921), under the implicit assumption that mates are random representatives of the population. In the classical one-locus Hardy–Weinberg model with only additive and dominance effects, this connection requires random pairing between female and male reproductive pools and no selection or other processes that change their allele frequencies from the reference population. With Mendelian segregation, the progeny then have the same allele frequencies and, consequently, the same Hardy– Weinberg genotype frequencies as the reference population. Under these conditions, the within-generation slope equals twice the parent-to-offspring slope. Consequently, additive genetic value was interpreted as an individual’s genetic merit as a parent and hence equivalent to breeding value, explaining why the terms are commonly treated as synonymous (Fisher 1919; Lush 1943; Falconer 1981; Lynch and Walsh 1998).

When the female and male reproductive pools differ, how-ever, a candidate’s expected progeny performance depends on the opposite reproductive pool (Falconer 1985). In what follows,we refer to additive genetic value as the value calculated from the slope of the within-generation linear projection of genetic value on allele dosage. We refer to breeding value as the value calculated from twice the slope of the between-generation linear projection of expected offspring genetic value on parental allele dosage, relative to a specified mate population. This dependence on the mate population connects breeding value to GCA.

Although commonly used to quantify expected parental performance in crosses between populations, GCA can also be defined for reproductive groups within a single population (Sprague and Tatum 1942; Griffing 1956b,a; Vitezica *et al*. 2016). For example, when females and males form the two mate pools, the GCA of a male is equal to the difference between the expected genetic value of the male’s progeny (generated by mating the male with all females) and the expected genetic value of all progeny (generated by mating all males with all females). This definition is similar to the operational (progeny-based) definition of transmitting ability (and hence half the breeding value), but with the explicit definition of the mate pool.

Imprinting is another genetic effect that contributes to the variation of complex traits (Wolf *et al*. 2008; Hager *et al*. 2008; Lawson *et al*. 2013; O’Doherty *et al*. 2015), which was not considered in the initial quantitative genetic models (O’Brien and Wolf 2019). It is an epigenetic effect often mediated by DNA methylation and histone modifications established during the formation of gametes. This methylation process differs between the formation of male and female gametes (such as ova and sperm), leading to differences in gene expression between maternally and paternally inherited alleles in progeny (Reik *et al*. 2001). The molecular basis of this process is now well understood (Tucci *et al*. 2019; Triantaphyllopoulos *et al*. 2016) and multiple genes have been characterized as maternally or paternally expressed (https://www.geneimprint.com/site/genes-by-status; Jirtle 1997; Hofmeister *et al*. 2025). One of the prominent examples is the IGF2 gene, which has been characterized as paternally expressed in multiple mammals (DeChiara *et al*. 1991; Giannoukakis *et al*. 1993; Van Laere *et al*. 2003; Brabazon *et al*. 2022; Ferguson-Smith *et al*. 1991; Suzuki *et al*. 2005).

Hill and Keightley (1988) presented a quantitative genetic model with imprinting, assuming complete silencing of imprinted alleles. Spencer (2002) extended the model to general imprinting by introducing different genotypic values for maternal and paternal heterozygotes. The term maternal heterozygote indicates that the alternative allele of a heterozygote was inherited from its maternal side. With imprinting, an individual’s genetic value can depend on whether an allele was inherited from its mother or father, whereas its expected progeny value can depend on whether the individual is the mother or father of the progeny. De Koning *et al*. (2002) used an explicit imprinting effect to describe the difference between maternal and paternal heterozygotes and studied the detection of imprinted QTL. Álvarez-Castro (2014) extended the Natural and Orthogonal Interactions (NOIA) framework to imprinting, distinguishing genetic effects defined relative to individual genotypes from those defined for a reference population. Schaeffer *et al*. (1989) formulated pedigree-based prediction models considering imprinting effects to estimate gametic variances and predict gametic values. Nishio and Satoh (2015) developed genomic prediction models with imprinting effects to estimate genetic variances and predict genomic values. O’Brien and Wolf (2019) derived several quantitative genetic parameters under the orthogonal model, including breeding values, non-additive genetic deviations, and their variances. These earlier derivations assumed random mating, except those of Álvarez-Castro (2014), which allowed for more general genotype frequencies. Recently, Krätschmer and Robinson (2026) analyzed a quantitative genetic model with indirect and parent-of-origin effects under random and assortative mating. Here, we consider random pairing between female and male mate pools that can differ in allele frequency, with and without imprinting.

### Theory

In the following, we work with the quantitative genetic model with additive, dominance, and imprinting effects (Álvarez-Castro 2014; Nishio and Satoh 2015; O’Brien and Wolf 2019; De Koning et al. 2002). Consider a bi-allelic locus with an alternative allele 1 of frequency *p* and reference allele 0 of frequency *q* = 1 *−p* with additive (*a*), dominance (*d*), and imprinting (*i*) effects. If *i* = 0 the model is equivalent to the standard additive and dominance model (Falconer 1981). At this locus, the genetic value of individual (*G*) with genotype 0*/*0 is*− a*, genotype 0*/*1 is *d* + *i*, genotype 1*/*0 is *d− i*, and genotype 1*/*1 is *a*. We write ordered genotypes with the maternally inherited allele first and the paternally inherited allele second.

Throughout the derivations, pairing is random between the female and male pools, and segregation is Mendelian. The reference population is used to define additive genetic value; the mate pool is used to define expected progeny genetic value and breeding value. Let *p*_*f*_ and *p*_*m*_ denote the allele frequencies respectively in the female and male mate pools, while *p* denotes the allele frequency in the whole reference population. We first consider the classical benchmark with undifferentiated mate pools, in which *p*_*f*_ = *p*_*m≠*_*p*, with Hardy–Weinberg genotype frequencies. This is equivalent to assuming random mating within the population. We then allow the female and male pools to differ in allele frequency (*p*_*f*_ = *p*_*m*_). In this differentiated mate-pool setting, pairing remains random between the pools. Finally, we extend both settings with imprinting effects.

### Standard quantitative genetic model

Fisher studied the variation of phenotypic values and decomposed the phenotypic value of an individual (*P*) into genetic value (*G*) and environmental deviation (*E*) (Fisher 1919, 1930, 1958). He further decomposed the genetic value of an individual into an additive genetic value and a non-additive genetic deviation by regressing the genetic values of individuals on their number of 0 and 1 alleles:

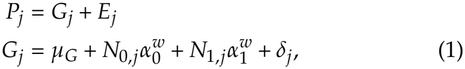

where *G*_*j*_ is the genetic value of an individual *j, µ*_*G*_ is the mean genetic value in the reference population, *N*_0,*j*_ and *N*_1,*j*_ are respectively the number of 0 and 1 alleles (dosage), 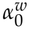 and 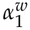 are allele effects, and *δ*_*j*_ is the residual. Here the allele effects are centered so that 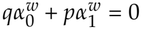. We use the superscript *w* to denote that these allele effects are defined within a generation (they characterize the decomposition of genetic values in the current population).

Since *N*_0,*j*_ + *N*_1,*j*_ = 2, we can express the same model using the within-generation slope 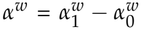 and centered allele dosage 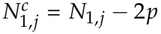, where 2*p* is the mean allele dosage in the reference population:

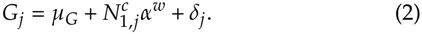

For a fixed reference population, centering the dosage changes the intercept but leaves the regression slope and fitted genetic values unchanged. The centering condition gives 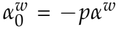 and 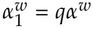. The within-generation slope *α*^*w*^ in (2) is defined for a reference population with non-zero allele dosage variance as:

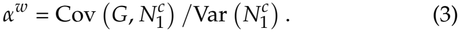

This slope is Fisher’s “average effect” of gene (allele) substitution (Fisher 1941; Falconer 1985). Using the decomposition (2), Fisher (1919, 1930, 1958) defined 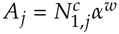 as additive genetic value and *δ*_*j*_ as non-additive genetic deviation. Note that these terms are part of (2).

**Under Hardy–Weinberg genotype frequencies from random mating within a population**, the expression for the slope *α*^*w*^ simplifies to (Fisher 1930, 1958; Falconer 1981, 1985):

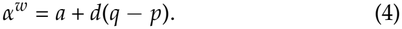

For other genotype frequencies, the slope is given by the general regression expression (3). In either case, *A*_*j*_ is defined by the within-generation regression of genetic value on allele dosage in the reference population. The Appendix A shows how this definition holds irrespective of the centering and how it connects to the widely used definitions in Falconer (1981).

Building on statistical foundations of quantitative genetics laid by Fisher (1919) and Wright (1921), Lush (1935, 1943) developed the quantitative-genetic treatment of breeding value, *B*_*j*_. He approached breeding value through progeny testing, expressing it as twice the difference between an individual’s expected progeny genetic value and the corresponding population progeny mean, with mates drawn randomly from the population. This definition is between generations, as it concerns the transmission of alleles, not genotypes, to offspring. Fisher’s additive genetic value, in contrast, is defined by the decomposition of genetic values of different genotypes within the reference population. Under the classical benchmark described above and without imprinting, breeding value equals additive genetic value. Although this equivalence depends on the mating conditions (Falconer 1985), breeding value and additive genetic value are often used interchangeably.

**With differentiated mate pools**, the additive genetic value and breeding value can differ (Fisher 1958, 1930; Falconer 1985), because the relationship between parental genotype and off-spring genetic value depends on the genotype distribution of both the selection candidates and their mates. In this case, let *b* denote the parent-to-offspring slope of the expected genetic value of the progeny of evaluated individuals E *G*_progeny_ on their centered allele dosage 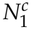. We call *α*^*b*^ = 2*b* the between-generation coefficient, on the conventional breeding-value scale:

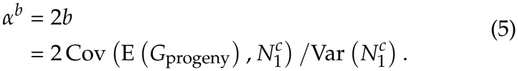

Here, E(*G*_progeny_) depends on the genotype of the evaluated individuals and the genotype of their mating candidates (each transmitting half of their genome). Appendix B illustrates the derivation of *α*^*b*^.

Based on this between-generation coefficient, the breeding value of the evaluated individuals is then 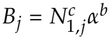 (e.g., Fisher 1958, 1930; Falconer 1985). This definition makes explicit that breeding value is a function of expected transmission to off-spring, rather than a component of the decomposition of *G*_*j*_ within the current generation. With undifferentiated mate pools and Hardy–Weinberg genotype frequencies, and with no parent-of-origin effect, expressions (3), (4), and (5) all give the same value, and the additive genetic value *A*_*j*_ equals the breeding value *B*_*j*_.

For females evaluated as parents, the mate pool consists of males. The between-generation coefficient for females 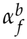 is then:

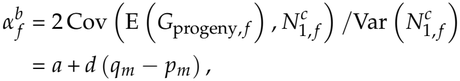

and for males 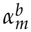 is:

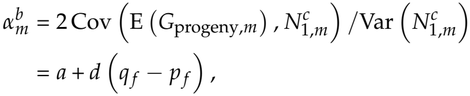

where E(*G*_progeny,*s*_) is the expected genetic value of progeny of the evaluated sex when mated with the other sex, 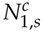 is the centered number of 1 alleles of the evaluated sex, and *p*_*s*_ and *q*_*s*_ are the allele frequencies of the opposite sex. Appendix B details the derivation for 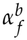 and 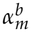. With undifferentiated mate 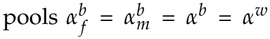. In general, the breeding value of an individual of sex *s* = {*f, m*} is 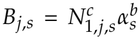, whereas the additive genetic value of an individual is 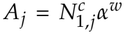, which can differ from the breeding value, and does not depend on the individual’s sex when the same reference population is used.

The above derivation highlights a fundamental distinction: additive genetic value is a within-generation quantity arising from the linear projection of genetic value of individuals on their allele dosage, whereas breeding value is a between-generation quantity arising from the linear projection of expected genetic value of progeny on parental allele dosage for a specified mate pool. Thus, breeding value is not a component of the decomposition of an individual’s genetic value in model (2). Appendix D uses the Price equation to connect these breeding-value expressions to the change in mean genetic value under selection (Price 1970, 1972). Specifically, this formulation identifies how the between-generation coefficients 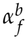 and 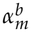 from the progeny regressions above enter the expression for response to selection.

Breeding value and GCA both refer to expected progeny genetic value, but use different scales (Griffing 1956b,a). GCA is the difference between a parent’s expected progeny value with a specified tester or mate population and the mean progeny value of all parents evaluated with that population (see also Falconer (1981) and Vitezica *et al*. (2016)). Indeed, GCA equals half the breeding value when both are defined relative to the same mean progeny value and expected progeny value is linear in parental allele dosage. This half-breeding-value quantity is also the transmitting ability used to predict genetic merit for the specified mate population and parental role (Yapp 1925; Beef Improvement Federation 1972; Falconer 1981). Figure 1 summarizes the within-generation definition of additive genetic value and the between-generation definition of breeding value, and shows its relationship to GCA.

**Figure 1.**
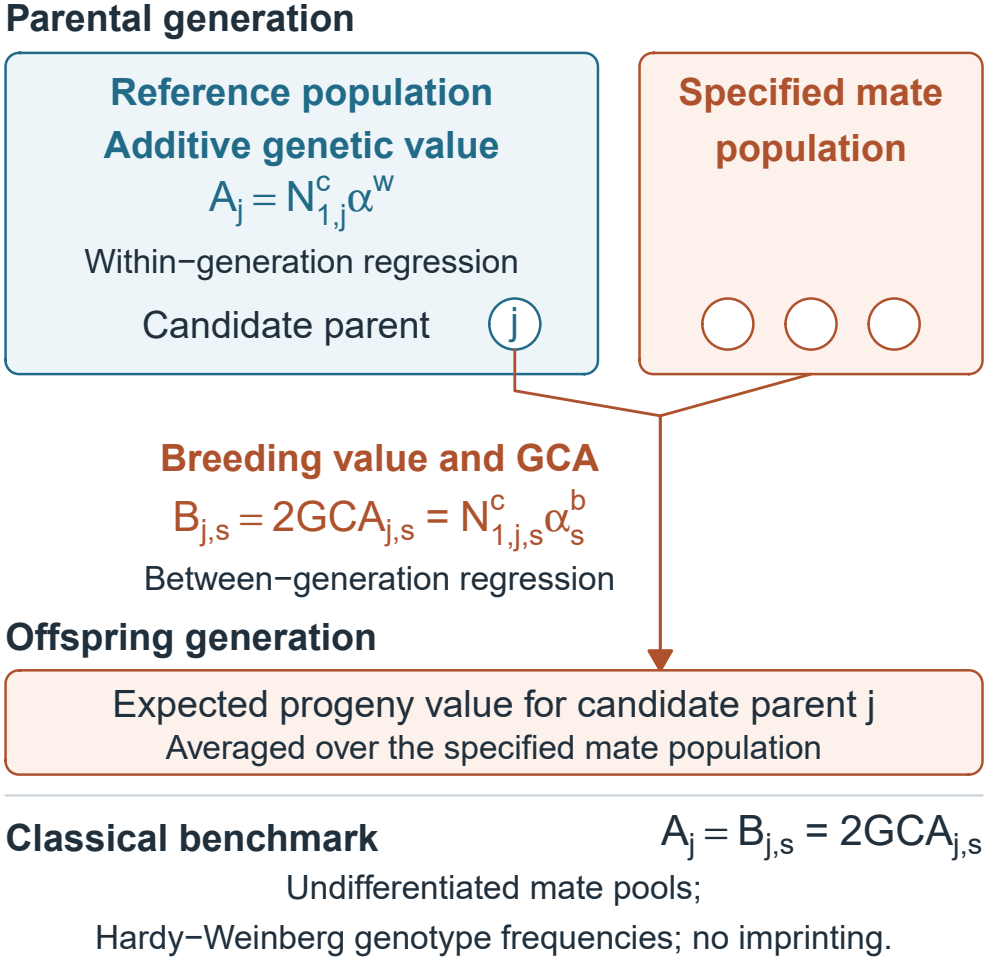
Additive genetic value, breeding value, and general combining ability (GCA) in a one-locus model with additive and dominance effects. Additive genetic value *A*_*j*_ is defined by the within-generation regression in the reference population. Breeding value *B*_*j,s*_ is defined from expected progeny genetic value under random pairing with the specified mate population, where *s* denotes the female or male parental role. General combining ability GCA_*j,s*_ = *B*_*j,s*_*/*2 uses the same mate popula-tion and progeny-mean baseline. 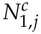 is allele dosage centered within the reference population, 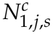 is allele dosage centered within the evaluated parental pool, *α*^*w*^ is the within-generation slope, and 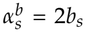 is twice the between-generation slope for parental role *s*.

### Quantitative genetic model with imprinting

With imprinting, the reciprocal heterozygotes 0*/*1 and 1*/*0 have respective genetic values of *d* + *i* and *d −i*. Consequently, the breeding values of males and females with the same ordered genotype usually differ, because their heterozygous progeny have different genetic values (O’Brien and Wolf 2019; Tier and Sölkner 1993; Neugebauer *et al*. 2010a,b). We first consider undifferentiated mate pools, then distinguish the reference used for additive genetic value from the differentiated mate pools used for breeding value.

Extending Fisher’s standard model (1,2) with imprinting (parent-of-origin) effects at bi-allelic loci gives (Spencer 2002; Schaeffer *et al*. 1989):

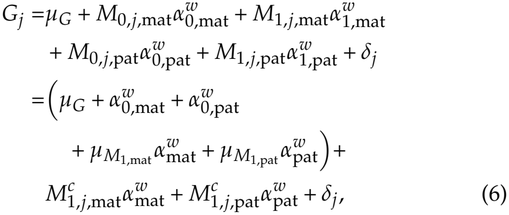

where *M*_*x,j,o*_ is the origin-specific allele dosage: the number of copies (0 or 1) of allele *x* inherited by individual *j* from its mother (*o* = mat) or father (*o* = pat). The total allele dosage then satisfies *N*_1,*j*_ = *M*_1,*j*,mat_ + *M*_1,*j*,pat_. The centered origin-specific allele dosage 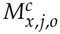 is obtained by subtracting the corresponding allele frequency in the reference population. Parental origin of alleles in heterozygotes must be known for this model. The within-generation slopes 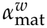 and 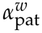 in (6) are defined jointly by least squares. When the reference population is formed by union of gametes under random mating, the two origin-specific allele dosages are uncorrelated, so the slopes can be written as:

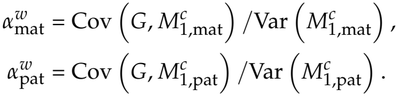

**With undifferentiated mate pools**, expressions for the slopes 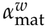 and 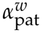 simplify to (O’Brien and Wolf 2019):

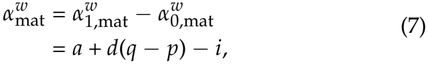

and

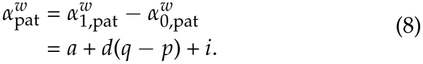

This gives the allele effects:

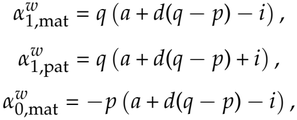

and

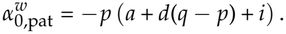

For given *a, d*, and *i*, these allele effects depend on allele frequency, while calculating an individual’s additive genetic value still requires parent-of-origin information.

Equation (6) defines the additive genetic value of individual *j* as 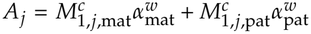 The breeding value of the same individual, evaluated as a female (*s* = *f*) or male (*s* = *m*) parent, is 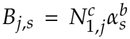.Here, mat and pat identify alleles in-herited from the individual’s own mother and father, whereas *f* and *m* on 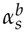 identify its role as a female or male parent of the next generation. With undifferentiated mate pools and ran-dom union of gametes 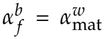 and 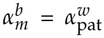. Even then, additive genetic value weights the two inherited alleles by their origins, while breeding value weights the total allele dosage by the individual’s parental role.

**With differentiated mate pools**, the female and male parental allele frequencies differ (*p*_*f*_ ≠ *p*_*m*_). Additive genetic value remains defined in the specified reference population by (6). Holding that reference fixed, changing the mate pools does not change the within-generation slopes or the centering of origin-specific allele dosages. However, for breeding value, expected progeny genetic value is averaged over the opposite-sex mate pool. Thus, the coefficient for a female parent depends on *p*_*m*_, and that for a male parent depends on *p*_*f*_. Applying (5) gives, for females:

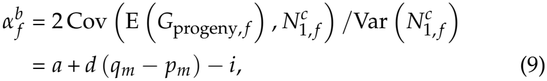

and for males 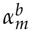 is:

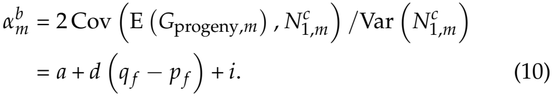

Therefore, the breeding value of an individual in parental pool *s* is 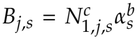, where 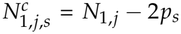 is centered within that parental pool.

Finally, to connect with the GCA, consider the progeny of female parent *j* and male parent *k*. Their expected genetic value can be written as:

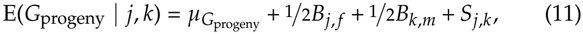

where 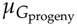 is the mean progeny genetic value across all possible pairings of female and male parental pools, *B*_*j, f*_ and *B*_*k,m*_ are the breeding values of the parents, and *S*_*j,k*_ is a residual. For a given parent *p* of sex *s*, averaging over the mate’s pool gives an expected progeny value of 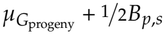 so their GCA is GCA_*p,s*_ = ½*B*_*p,s*_, and the residual *S* represents the specific combining ability (SCA) of the parental pair. An individual offspring can additionally deviate from E(*G*_progeny_ |*j, k*) due to segregation.

### Demonstration

To illustrate the difference between the within-generation 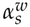 and the between-generation 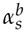, we compare them under undifferentiated and differentiated mate pools, without and with imprinting. Initially, we assume a single bi-allelic locus with alleles 0 and 1, additive effect *a* = 1, dominance effect *d* = 0.75, no imprinting effect (*i* = 0), equal proportion of females and males, and allele frequency in females of *p*_*f*_ = 0.3 and in males *p*_*m*_ = 0.9. We assume Hardy–Weinberg genotype frequencies within females and males. We calculate the within-generation 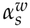 separately for females at *p*_*f*_ = 0.3 and males at *p*_*m*_ = 0.9. For the between-generation 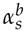, we assume random pairing between females and males. For comparison, we also calculate both quantities under the classical benchmark, where the reference population has Hardy–Weinberg genotype frequencies at *p* = 0.6, and the mate pools are undifferentiated (*p*_*f*_ = *p*_*m*_ = *p*). These results are presented in Figure 2. The corresponding additive genetic values and breeding values derived from these quantities are shown and discussed in Appendix C (Table C1).

**Figure 2.**
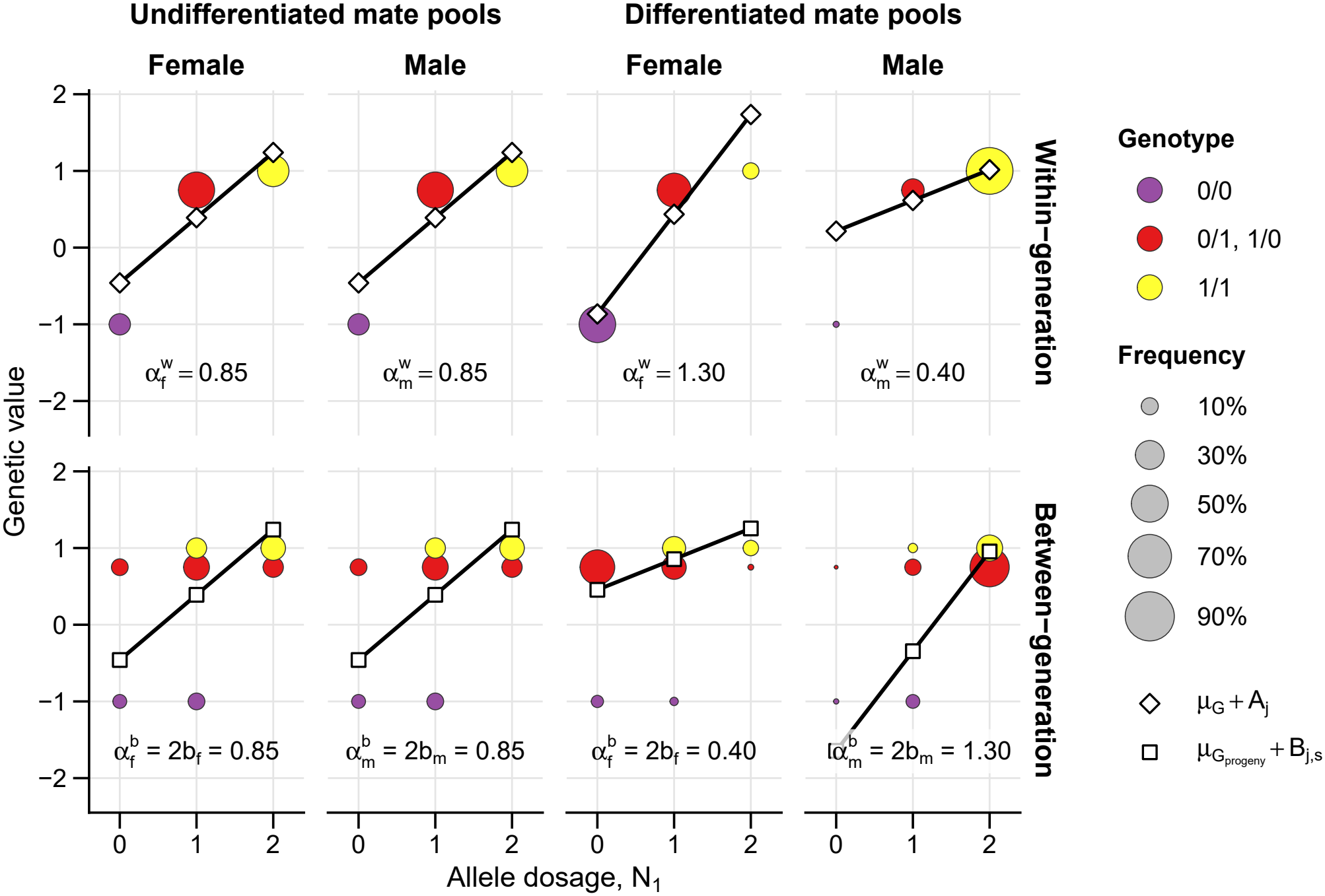
Within- and between-generation regressions in undifferentiated and differentiated mate pools, without imprinting. The trait is controlled by a single bi-allelic locus with additive effect *a* = 1, dominance effect *d* = 0.75, and no imprinting effect. Undifferentiated mate pools have *p*_*f*_ = *p*_*m*_ = *p* = 0.6; while differentiated female and male pools have *p*_*f*_ = 0.3 and *p*_*m*_ = 0.9. Reference populations have Hardy–Weinberg genotype frequencies. Bubbles denote genetic values and colors identify genotypes. In the top row, bubbles show individual genetic values *G*_*j*_ against allele dosage *N*_1_. Their area denotes genotype frequency in the female or male reference population. Diamonds and the solid line show fitted genetic value *µ*_*G*_ + *A*_*j*_, with slope 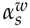, giving additive genetic value *A*_*j*_. In the bottom row, bubbles show possible progeny genetic values against parental allele dosage *N*_1_. Their area denotes the frequency of each parent–progeny combination under random pairing. Squares and the solid line show fitted genetic value 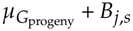. with slope 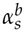. giving breeding value *B*_*j,s*_. See Table C1 for the actual additive genetic values and breeding values.

Under the classical benchmark, the within-generation 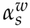 for females and males are identical (0.85), when calculated for the Hardy–Weinberg reference population at *p* = 0.6 (Figure 2, top-left). Under separate female and male reference populations, however, the within-generation 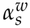 for females (1.30) is larger than for males (0.40) (Figure 2, top-right). The mean of the two slopes is 1.30 *×* 0.5 + 0.40*×* 0.5 = 0.85, the slope in the classical benchmark. This equality follows because *a* + *d*(*q −p*) is linear in *p*, but does not give the regression slope in the pooled genotype distribution.

The between-generation 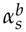 under the classical benchmark is identical across roles (0.85) and coincides with 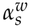 (Figure 2,bottom-left). This equivalence arises because the evaluated individuals and their mating partners have the same Hardy– Weinberg genotype frequencies in the classical benchmark. Random pairing produces these frequencies also in their progeny, so all three groups share the same allele and genotype frequencies (Fisher 1930, 1958; Falconer 1985). Finally, under differentiated female and male mate pools, 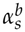 for females (0.40) is smaller than for males (1.30) (Figure 2, bottom-right). In this example,the between-generation 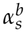 equals the within-generation 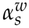 in the opposite-sex reference population, because each parent’s mate pool consists of individuals of the opposite sex. The large difference between the within-generation 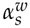 and the between-generation 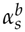 in the same sex (1.30 vs. 0.40 in females) reflects the distinction between a within-generation statistical decomposition and a between-generation transmission process.

The above results are well known and have been used in genomic prediction for crossbreeding (Ibáñez-Escriche et al. 2009; Hayes et al. 2009; Vitezica et al. 2016). The different between-generation 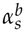 show that females and males with the same genotype can have different breeding values. See Table C1 in Appendix C, to compare breeding values for individual genotypes between sexes. This difference in 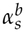 and corresponding breeding values arises due to dominance and different allele frequencies in the female and male mate pools, even without imprinting.

We repeat the analysis with an imprinting effect *i* = 0.5, retaining the other assumptions. Figure 3 shows the within-generation and between-generation analysis, while additive genetic values and breeding values are shown and discussed in Appendix C (Table C2). Under imprinting, the decomposition of genetic value yields two distinct regression coefficients, one for each parent of origin ((6)). In Figure 3 we illustrate how these within-generation regressions are calculated from the genetic values of ordered genotypes. Because these regressions operate at the haploid gamete level, the origin-specific allele dosages range from 0 to 1. This contrasts with the between-generation regressions and Figure 2 regressions, which operate at the diploid genotype level where allele dosages range from 0 to 2. Figure C1 combines parent and progeny genetic values, the origin-specific within-generation (gamete) regressions and resulting additive genetic values, with the between-generation (genotype) regressions and resulting breeding values.

**Figure 3.**
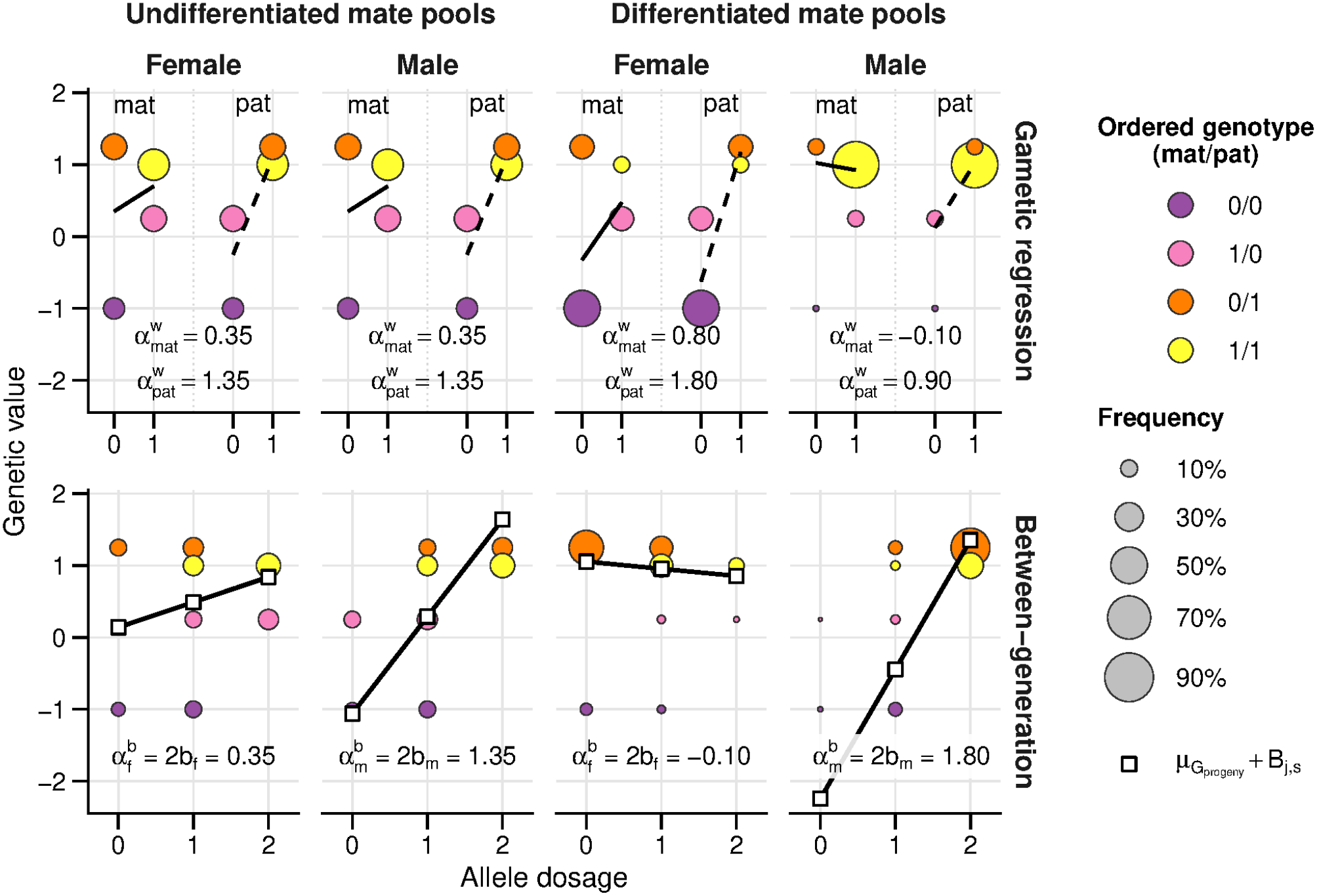
Within- and between-generation regressions in undifferentiated and differentiated mate pools, with imprinting. The trait is controlled by a single bi-allelic locus with additive effect *a* = 1, dominance effect *d* = 0.75, and imprinting effect *i* = 0.5. Undifferentiated mate pools have *p*_*f*_ = *p*_*m*_ = *p* = 0.6, while differentiated female and male pools have *p*_*f*_ = 0.3 and *p*_*m*_ = 0.9. Reference populations have Hardy–Weinberg genotype frequencies. Bubbles denote genetic values and colors identify ordered genotypes. In the top row, bubbles show individual genetic values *G*_*j*_ against origin-specific allele dosage *M*_1_ and recognizing the parental origin of alleles. Their area denotes genotype frequency in the female or male reference population. Solid and dashed lines show within-generation gamete regressions respectively giving 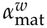 and 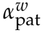. In the bottom row, bubbles show possible progeny genetic values against parental allele dosage *N*_1_. Their area denotes the frequency of each parent–progeny combination under random pairing. Squares and the solid line show fitted genetic value 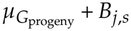, with slope 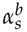 giving breeding value *B*_*j,s*_. See Table C2 for the actual additive genetic values and breeding values.

For the common reference population with allele frequency *p* = 0.6, we obtain the within-generation 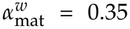 and 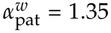 for both females and males (Figure 3, top-left). For the separate female reference population at *p*_*f*_ = 0.3, we obtain 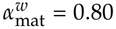 and 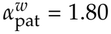.. For the male reference population at *p*_*m*_ = 0.9, we obtain 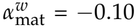 and 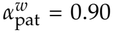 (Figure 3,top-right). Each reference population has both maternally and paternally inherited alleles. For individuals evaluated as female and male parents under undifferentiated mate pools, we ob-tain the between-generation 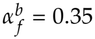 and 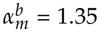 (Figure 3, bottom-left), respectively equal to the within-generation 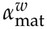 and 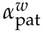. With differentiated mate pools, the corresponding values are 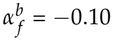 and 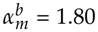 (Figure 3, bottom-right).

To the best of our knowledge, these gamete-level results have not been previously presented or interpreted in this manner. With undifferentiated mate pools, each origin-specific within-generation coefficient is the same in the female and male reference populations, although the maternal and paternal coefficients differ under imprinting. These coefficients equal the between-generation coefficients for the corresponding transmit-ting parental roles 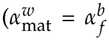 and 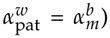. This condition is met here because our example assumes that the genotype frequencies within pools follow Hardy–Weinberg equilibrium. Consistent with the theory, these results demonstrate that under imprinting, females and males with identical genotypes can have different breeding values. See Table C2 in Appendix C to compare breeding values for individual genotypes between sexes.

With differentiated mate pools, the within-generation coef-ficients (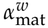 and 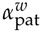) differ between the female and male ref-erence populations. In this example, 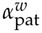 evaluated within the female pool equals 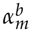, while 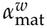 evaluated within the male pool equals 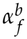. This symmetry means that the change in additive genetic value due to inheriting a different paternal allele into a female pool is equal to the change in breeding value due to transmitting a different allele from a male.

The above example with imprinting is for a specific set of allele frequencies in males and females. Figure 4 shows the difference between the between-generation 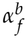 for females and 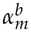 for males, defined as 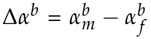, across a wider spectrum of allele frequencies, with additive effect *a* = 1, dominance effects *d ∈ {*0, 0.5, 1, 1.5*}*, and imprinting effects *i ∈ {−*0.5, 0, 0.5*}*. From Equations (9) and (10), Δ*α*^*b*^ = 2*d*(*p*_*m*_ *− p*_*f*_) + 2*i*. When there are no dominance and imprinting effects (*d* = *i* = 0), Δ*α*^*b*^ = 0, be-cause the genetic values are a function of only the additive effect (*a*). In this case, both the within-generation 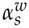 and between-generation 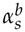 have the same value, and this value is the same for each sex. When *d≠*0 and *i* = 0, Δ*α*^*b*^ = 2*d*(*p*_*m*_*− p*_*f*_), which changes linearly with the allele-frequency difference between sexes. If the imprinting effect has a non-zero value and the dominance effect is 0, then Δ*α*^*b*^ = 2*i* holds across all allele-frequency combinations within the same scenario. For *d ≠* 0, Δ*α*^*b*^ = 0 when *p*_*m*_ *− p*_*f*_ = *−i/d*.

**Figure 4.**
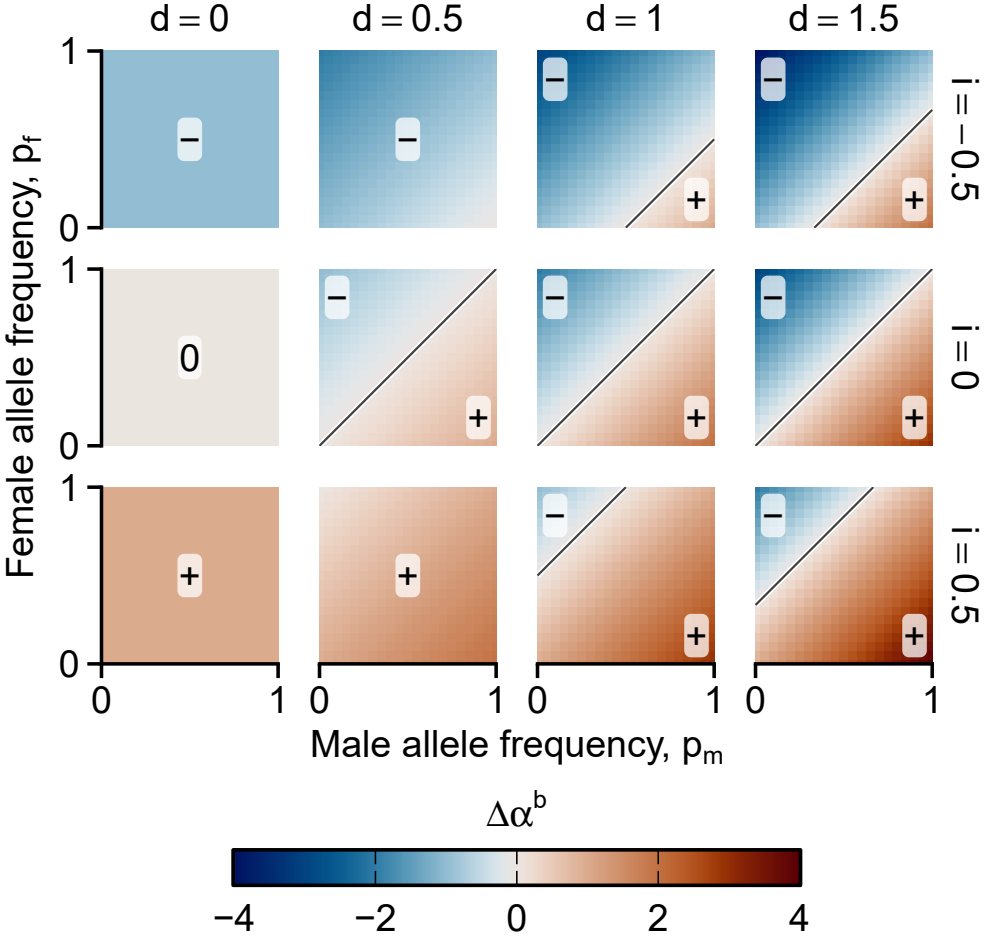
Difference in between-generation 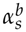 for male and female parents, 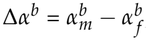, under differentiated mate pools at a locus as a function of allele frequency in females and males and genetic effects. Additive effect *a* = 1, dominance effects *d∈{* 0, 0.5, 1, 1.5}, and imprinting effects *i* ∈ {−0.5, 0, 0.5}. Lines mark zero differences; signs indicate direction of difference.

### Implications

The theory shows that additive genetic value and breeding value can differ with dominance and differentiated mate pools, or with imprinting. Reproductive groups can differ in allele frequency because of sex-specific (pre-)selection or recruitment from genetically differentiated populations. We focus on two selection scenarios that can generate these differences: antagonistic selection (Lucotte et al. 2016) and synergistic selection in the presence of parent-of-origin effects (O’Brien and Wolf 2019). We illustrate these scenarios by simulation and examine allele frequency differences in a population of Rubia Gallega beef cattle.

#### Two sources of allele frequency differences between sexes: A simulation

To illustrate how antagonistic and synergistic selection can drive differences in allele frequencies between the sexes, we used the AlphaSimR simulation package (Gaynor et al. 2021) with an extension for imprinting effects. We started the simulation with a base population of 2000 individuals, 1000 males and 1000 females. The simulated genome had 10 chromosomes with 1000 segregating sites per chromosome, with an expected alternative allele frequency of 0.5 at each site in each sex. Each site had an additive effect (*a*) and an imprinting effect (*i*), with no dominance effect (*d* = 0). To generate some large and many small effects, additive effects were generated using the gamma option in AlphaSimR, with shape parameter 0.001 and target additive variance of 1 in the base population. We used the same set of absolute effect sizes for additive and imprinting effects. For imprinting, these sizes were randomly assigned to loci and given positive or negative signs with equal probability. Figure 5 shows the additive and imprinting effects at each site.

**Figure 5.**
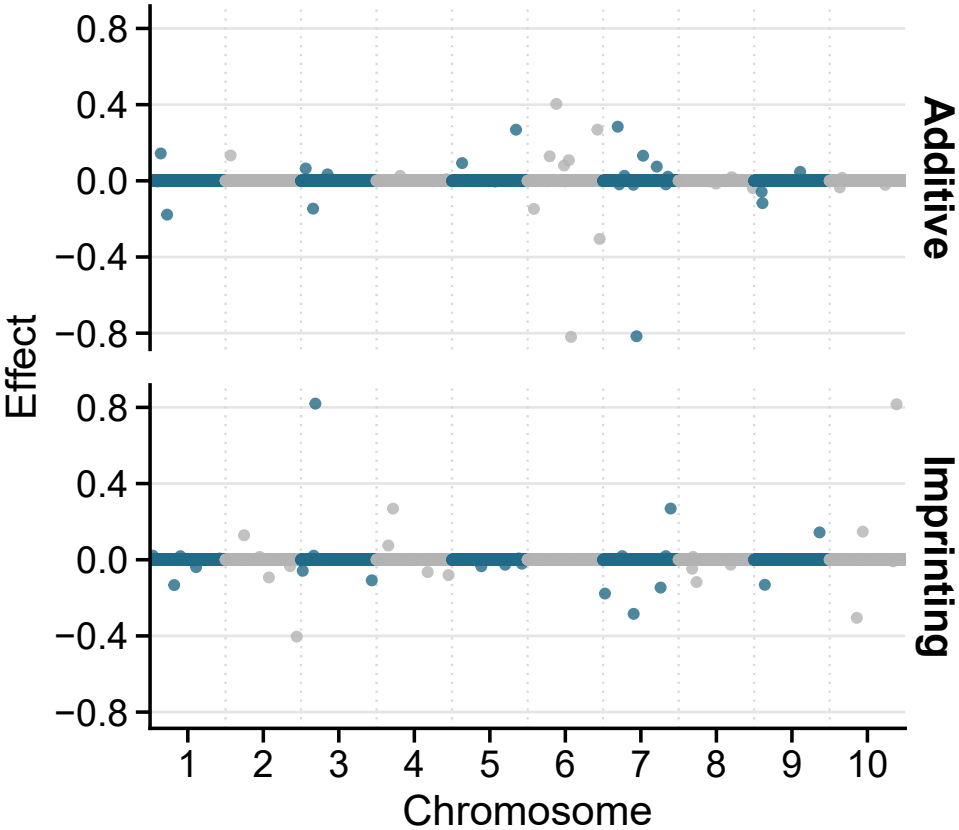
Additive (*a*) effects (top) and imprinting (*i*) effects (bottom) in the simulation. Each point represents a segregating site. Alternating blue and gray distinguish adjacent chromosomes.

We applied two selection strategies to the same base pop-ulation, using true genetic values or true breeding values as specified below:

1. Antagonistic selection: the 10% of females with the lowest genetic value (*G*_*j*_) and the 10% of males with the highest genetic value *G*_*j*_ were selected as parents.
2. Synergistic selection: the 10% of each sex with the highest breeding value for their parental role (*B*_*j,s*_) were selected as parents.

After one round of parental selection, we calculated the absolute allele frequency difference |*p*_*f*_ *−p*_*m*_| at each site between the selected female and male parental pools (Figure 6). Under antagonistic selection, larger absolute allele frequency differences were associated with larger absolute additive effects (Figure 6, top-left). There was no clear increase in these differences with absolute imprinting effect (Figure 6, bottom-left). Under synergistic selection, larger absolute differences instead occurred at sites with large absolute imprinting effects (Figure 6, bottomright). There was no clear increase with absolute additive effect (Figure 6, top-right).

**Figure 6.**
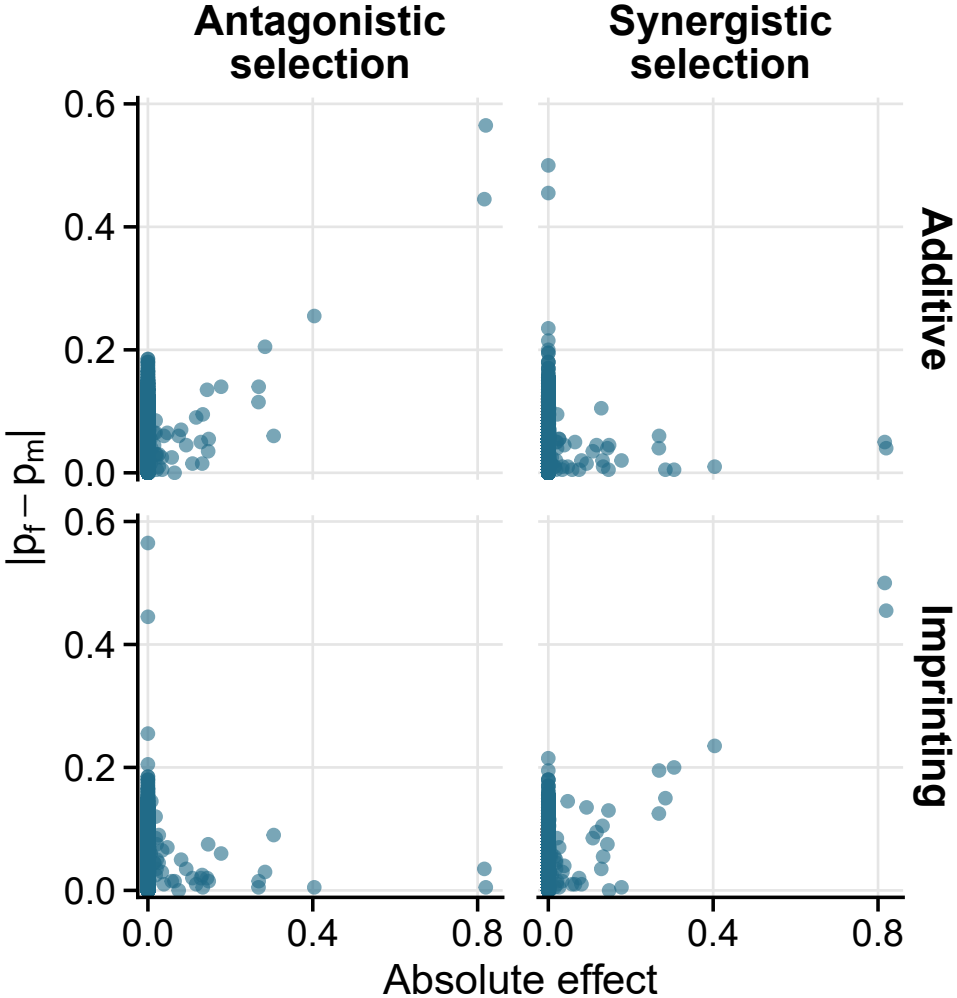
Absolute allele frequency difference |*p*_*f*_ *−p*_*m*_| across segregating sites between selected female and male parental pools in relation to absolute additive effect |*a* | (top row) and absolute imprinting effect |*i*| (bottom row). Columns show antagonistic selection (left) and synergistic selection (right).

These patterns are consistent with the regressions used to define additive genetic value and breeding value. In the expected base-population distribution, both origin-specific allele frequencies are 0.5. With no dominance, the within-generation slopes are 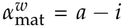 and 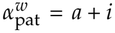. The imprinting terms cancel in the regression of genetic value on total allele dosage, whose slope is *a*, consistent with the association with additive effects under antagonistic selection. For selection on breeding value,the between-generation coefficients are 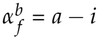 and 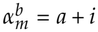 (see (9) and (10)). Imprinting therefore contributes with opposite signs to the coefficients for the two parental roles, so selection for higher breeding value can favor different alleles in female and male parental pools.

### Allele frequency differences between sexes: A real population

To examine possible allele frequency differences between female and male reproductive groups, we analyzed SNP marker genotypes from 4,531 Rubia Gallega beef cattle (3,631 females and 900 males). Individuals were genotyped using the Axiom BovMDv3 array (Thermo Fisher Scientific, Waltham, MA, USA). Genotypes were filtered using PLINK v1.9 (Purcell et al. 2007), with a minimum minor allele frequency of 0.01 and maximum missing rates of 0.10 per SNP and per individual. After filtering, 45,299 SNP markers remained.

We calculated the absolute allele frequency difference |*p*_*f*_ *−p*_*m*_ | for each SNP and averaged these differences within slid-ing windows of 50 SNP markers (Figure 7). Window means ranged from approximately 0.0% to 4.4%. We tested at a marker level which mean absolute allele frequency differences were significantly different from zero using Fisher’s exact test (Fisher 1922) and Bonferroni correction (Bonferroni 1936). Figure 7 high-lights four genomic regions: BTA2 (747,465-8,824,176 bp), BTA4 (114,416,811-114,439,054 bp), BTA6 (70,680,628-71,749,458 bp), and BTA17 (74,723,634-74,925,889 bp).

**Figure 7.**
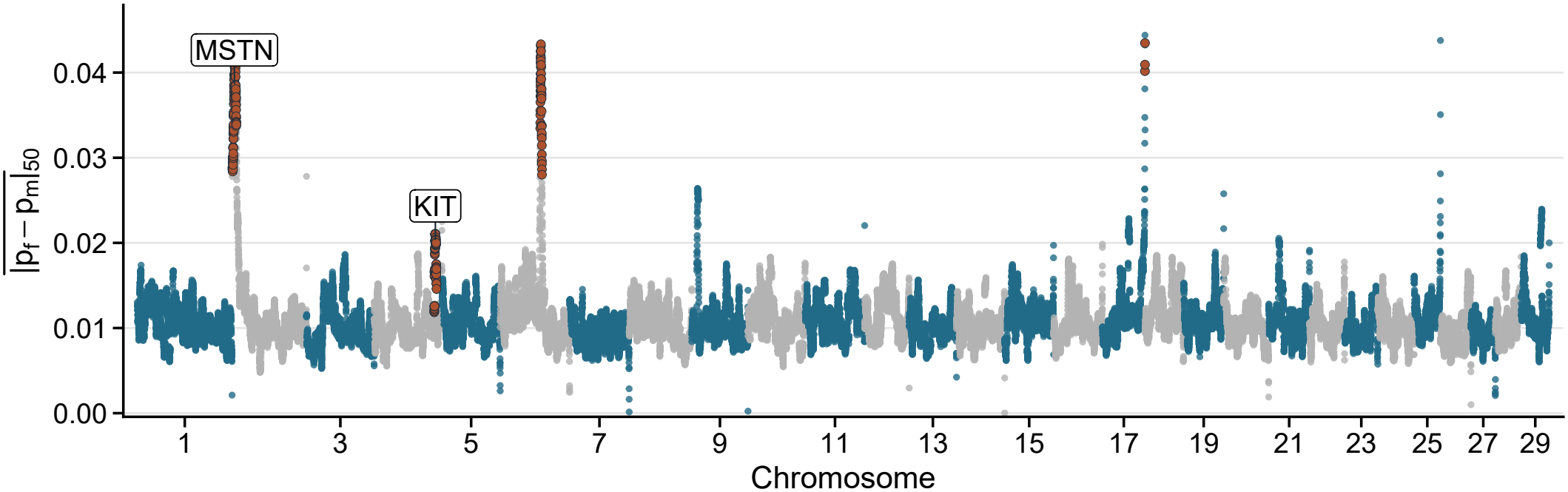
Mean absolute allele frequency difference between female and male reproductive groups in Rubia Gallega beef cattle,shown across chromosomes. Each point represents a mean over a sliding window of 50 SNP markers, 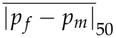. Alternating blue and gray distinguish adjacent chromosomes. orange outlined points mark the four regions discussed in the text.

## Discussion

This study reviewed the quantitative genetic theory of breeding value in a model with additive and dominance effects under undifferentiated and differentiated mate pools, and then extended this theory with imprinting effects. The review highlighted how breeding values depend on the specified mate population and its allele frequencies (Fisher 1930, 1958; Falconer 1985). In the model without imprinting, the within-generation *α*^*w*^ describes the linear regression of individual’s genetic value on allele dosage in the reference population, whereas the between-generation *α*^*b*^ is twice the slope of the linear regression of expected progeny genetic value on parental allele dosage for a specified mate pool. These respectively give additive genetic value *A*_*j*_ and breeding value *B*_*j,s*_. The Price equation treatment in Appendix D connects these breeding-value expressions to the change in mean genetic value under selection. With a common reference and mate population at Hardy–Weinberg genotype frequencies, and in the absence of imprinting, the two coefficients and values coincide, recovering the classical result (Fisher 1930, 1958; Falconer 1985).

These two coefficients can however differ with dominance and differentiated mate pools, or with imprinting, as well as other non-Mendelian inheritance patterns. Even with random pairing between female and male candidates, the mate pool can differ according to the individual’s role as a female or male par-ent. This distinction is relevant to selective breeding programs, which routinely use several reproductive groups within the population (males, females, young, old, pre-selected, etc.), each with a different set of expected mates and a different contribution to the next generation (Lush 1943; Antonios *et al*. 2023; Pardo *et al*. 2023; López-Carbonell *et al*. 2024). The population used to define additive genetic value therefore need not be the mate population used to define breeding value.

With imprinting, inherited parental origin affects the individual’s genetic value, while its role as a female or male parent affects expected progeny genetic value. A female and male with the same ordered genotype have the same additive genetic value in the same reference population, but can have different breeding values. Recent complementary theory examined how indirect and parent-of-origin effects alter genetic variance and resemblance among relatives under assortative mating (Krätschmer and Robinson 2026). Here, we focus on the expected progeny genetic value for a specified mate population and parental role, and its relationship to additive genetic value and GCA. In the models considered here, breeding value is twice GCA when both use the same mate population and mean progeny value (Griffing 1956b,a; Falconer 1981; Vitezica *et al*. 2016).

The most immediate application of this work is a setting where allele frequencies differ between sires and dams. Our simulation illustrates that such differences can arise from antagonistic or synergistic selection in line with the work of Lucotte *et al*. (2016). This simulation was inspired by the Rubia Gallega population, where we observed allele frequency differences between the active reproductive groups. This is not unexpected, because in beef cattle, sires and dams may be selected using different indexes (Roughsedge *et al*. 2005a,b; Twomey *et al*. 2020); sire selection often focuses on growth and conformation (Amer *et al*. 1998; Twomey *et al*. 2020), while dam selection gives more emphasis to maternal and reproductive traits (Roughsedge *et al*. 2005a; Twomey et al. 2020). Whether these objectives produce antagonistic selection at a locus depends on its effects on the selected traits.

In Rubia Gallega cattle, we observed prominent allele frequency differences between sires and dams in several genomic regions. The BTA2 region (747,465–8,824,176 bp) includes the Myostatin (MSTN) gene, which is linked to double muscling phenotype (Grobet et al. 1997; McPherron and Lee 1997; Kambadur et al. 1997). This region has also been associated with carcass traits in Rubia Gallega (Martinez-Castillero et al. 2021). Selection for double muscling on the male side and against calving difficulty on the female side could contribute to allele frequency differences in this region. On BTA6, earlier studies reported QTLs for birth weight, yearling weight, hot carcass weight, and *longissimus* muscle area (Davis et al. 1998; Casas *et al*. 2000), in the region of pronounced allele frequency differences (70,680,628–71,749,458 bp). These trait associations suggest possible selection-related explanations, but the present analysis does not distinguish selection from differences in relatedness, birth cohorts, or sampling between the reproductive groups.

Different selection criteria on male and female side also occur in dairy cattle. Fertility traits have received substantial attention in the last decades (Lucy 2001; Cole and VanRaden 2018), and fertility depends on both female and male contributions (Freeman 1984; Kropp et al. 2014; Jaton et al. 2017). In practice, dams are selected on a combination of female reproductive traits, while sires used in artificial insemination are also selected on semen production and quality (Norman et al. 2008; Shi et al. 2023). The extent to which these criteria and differences in selection intensity produce allele frequency differences between reproductive groups remains to be evaluated.

Allele frequency differences between sexes have also been reported beyond livestock. In a hybrid zone between the cyprinid fish *Barbus barbus* and *Barbus meridionalis*, Chenuil *et al*. (2004) considered differences in survival and fertility as possible explanations, alongside dispersal and cohort differences. Flintham *et al*. (2026) describe that allele frequency differences between reproductive males and females can arise due to sexually antagonistic selection. This process has been associated with differences in gene expression between sexes (Flintham 2025), which can be due to imprinting. Graffelman and Weir (2018) also reported differences at some markers in human samples from the 1000 Genomes Project.

Pre-selection is a plausible source of observed allele frequency differences between reproductive groups. In many animal breeding programs, artificial insemination sires, dams of sires, and regular dams have different roles in generating the next generation and are subject to different selection intensities and possibly also different selection indexes, either directly or indirectly. For example, artificial insemination sires are sons of the top sires and dams (first selection step), but are also selected based on additional testing (second selection step). Selection of these sires is more intense than that for their potential female mates. The definitions described in this study show how to calculate breeding values at each of these stages using the allele frequencies of the specified mate population. The reference population used in a genetic evaluation (Henderson 1984; Mrode and Pocrnic 2023) need not represent those intended mates. Specifying the mate population at each stage makes this distinction explicit.

Animal crossbreeding schemes, used in pigs, poultry, and rabbits, provide another setting where breeding values depend on the mate population. In these schemes, individuals from different purebred lines, say A, B, and C, are mated to produce AB and (AB)C crosses. Purebred individuals from A and B lines are usually genetically evaluated by a multi-trait analysis of purebred and crossbred performance (e.g. Wei and van der Werf 1994). Each population plays a specific role in such schemes, and the populations differ in allele frequencies. Imprinting effects, as described for the IGF2 gene in pigs (Van Laere *et al*. 2003), also require a distinction between reciprocal parental roles. Standard multi-trait analyses of purebred and crossbred performance do not automatically make this distinction. For example, A females may be evaluated for AB progeny with B males, while AB females are evaluated for (AB)C progeny with C males. The relevant breeding value in these scenarios depends on both the mate line and the parental role.

This dependence on the mate population is well known in hybrid plant breeding, where GCA is used to select individuals within each heterotic pool based on their hybrid progeny performance (Griffing 1956b,a). When imprinting affects progeny performance (Scott *et al*. 1998; Swanson-Wagner *et al*. 2009; Köhler *et al*. 2012; Castillo-Bravo *et al*. 2022), GCA can be defined separately for female and male parental roles. An individual can then have two GCA values, according to its role in the mating.

In the extreme case, these definitions can be leveraged to optimize mate allocation (e.g., Toro and Varona 2010; Werner et al. 2023) by evaluating single individuals as pools instead of groups of individuals. In our model, the expected value of a particular mating includes one-half of each parent’s breeding value and a specific combining ability term, in addition to the mean progeny genetic value (11). Breeding values alone therefore do not describe all differences between matings (crosses), which is well known in plant breeding (Griffing 1956a,b). Although we focused on expected progeny genetic value, mate allocation can also consider the genetic variance among progeny (e.g. Hohenboken 1985; Khaw et al. 2016; Yoshida et al. 2018; Sae-Lim *et al*. 2012).

We treated imprinting effects as fixed, but their biological basis can involve the dynamic process of DNA methylation (Reik et al. 2001; Triantaphyllopoulos et al. 2016; Tucci et al. 2019). Data on methylation state may help model variation in these effects beyond the fixed effects used here. Ayres *et al*. (2025) proposed an extension of the standard quantitative genetics model using methylation state data.

However, methylation states can vary across loci, tissues, sexes, time, and environmental conditions, and these sources of variation should be considered when using such data. Predicting progeny performance considering methylation state would then require a model of how methylation states change. The framework presented in this work could leverage such a model.

The distinction between additive genetic value and breeding value is also relevant to sex chromosomes, and mitochondrial and chloroplast DNA, which deviate from Mendelian transmission. These components of an individual’s genome can affect phenotype (e.g., Su et al. 2014; Khramtsova et al. 2019; Pacheco et al. 2020; Sanchez et al. 2023), but their transmission differs from that of the autosomal loci considered here. For example, Mafra Fortuna *et al*. (2024) defined sex-specific additive genetic values and breeding values with respect to mitochondrial DNA, because the phenotypes of both sexes can be affected by mitochondrial DNA. For X-linked loci in mammals, the model may also need to account for X inactivation. For both X- and Y-linked loci, breeding values must reflect sex-specific inheritance. Extending our approach to these cases requires the corresponding transmission model when calculating expected progeny genetic value.

In applications, genetic merit is estimated from observed phenotypes and pedigree or genotype data. Whether an analysis estimates additive genetic value or breeding value is determined by its statistical model and specified reference and mate populations, not by whether the information comes from an individual’s own or its progeny’s phenotypes. Both types of records can contribute to the estimation of different measures of genetic merit. In breeding programs, only some individuals have progeny, and these parents are usually pre-selected. Such selection and mate allocation can change the relevant mate distributions and whether the estimates are biased depends on how the statistical procedure accounts for these processes. Linear mixed models for genetic evaluation use pedigree or genotype information to combine phenotypic information within and between generations (Henderson 1975, 1984; VanRaden 2008; Aguilar et al. 2010; Meuwissen et al. 2001). Standard statistical procedures estimate additive effects defined by the fitted model, but they do not automatically distinguish breeding values for different mate populations or parental roles when differentiated mate pools or imprinting are present. GCA-type models offer one possible way forward (Griffing 1956a,b; de Jong *et al*. 2023), given the connection between GCA and general definition of breeding value in this work. Estimating such breeding values in the popular GBLUP procedure requires an explicitly defined covariance structure, which was not derived or tested in this study (Wei and van der Werf 1994; Christensen *et al*. 2014, 2015).

Another popular procedure is SNP-BLUP, where fitted marker effects describe genotype–phenotype associations under an assumed genetic model and reference population (Meuwissen *et al*. 2001). Assuming that the parametrization of the fitted model matches the one used in our one-locus expressions, estimates from this procedure could be used with our expressions to calculate breeding values for specified mate pools and their allele frequencies. In this context, the Natural and Orthogonal InterAction (NOIA) model (Alvarez-Castro and Carlborg 2007; Álvarez-Castro 2014) provides a particularly suitable parameterization of genetic effects. By maintaining orthogonality (i.e., uncorrelated effects) under arbitrary allele frequencies and departures from Hardy–Weinberg equilibrium, NOIA enables the simultaneous estimation of additive (*a*), dominance (*d*), and imprinting (*i*) effects. Under the NOIA framework, breeding values are not direct outputs of the statistical model. They are constructed quantities that depend jointly on the estimated genetic effects and the allele frequencies of potential mating candidates, as demonstrated in our theoretical derivations. How best to estimate these quantities jointly across loci requires further work.

Knowledge of genomic regions or genes associated with trait variation helps us understand the biological basis of traits. It can also be used to give different weights to loci in genomic prediction (Calus *et al*. 2014; Tiezzi and Maltecca 2015; Karaman *et al*. 2018; Wang *et al*. 2012; Zhang *et al*. 2010). Genome-wide association studies commonly estimate allele substitution effects (Uffelmann *et al*. 2021), but can be extended with dominance (e.g., Sun *et al*. 2014; Lopes *et al*. 2014; Vitezica *et al*. 2017) and imprinting terms (e.g., Cheng *et al*. 2013; Hu *et al*. 2015; Álvarez-Castro 2014; Jiang *et al*. 2017). Interpreting imprinting in such analyses requires distinguishing the genetic contrast between reciprocal heterozygotes from population-dependent allele substitution effects. In our model, this contrast is Δ*α*^*b*^ = (*d* + *i*) *−* (*d− i*) = 2*i*, independent of allele frequencies, whereas substitution effects and their associated genetic variances can depend on these frequencies (Falconer 1981; Lynch and Walsh 1998; Varona *et al*. 2018). For example, under random union of gametes, dominance and unequal allele frequencies among maternally and paternally inherited alleles can make 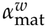 and 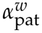 differ even without imprinting. Likewise, in progeny-based analyses, dominance and unequal allele frequencies between male and female mate pools can make 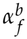 and 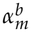 differ without imprinting (Figure 4). Such differences therefore cannot, on their own, be interpreted as evidence of imprinting.

Under the definition developed here, breeding values quantify the expected genetic contribution of selection candidates to the next generation and depend on the allele frequencies of their potential mates. This perspective focuses primarily on maximizing the response to selection in the subsequent generation. For this purpose, breeding values should be calculated for each generation using the allele frequencies of the intended mate pools. With dominance, these frequencies affect the between-generation coefficients, so breeding values can change between generations even when the underlying genotypic effects remain fixed.

However, selecting for the greatest short-term response need not maximize long-term genetic gain. For example, Wientjes *et al*. (2022) found that, in their simulations with epistasis, genomic selection gave a larger initial response than phenotypic selection, but a smaller total response after 50 generations, mainly because more genetic variation was lost. This illustrates the need to consider how selection for the next generation affects the opportunities for selection in later generations. For our framework, one possibility is to evaluate candidates for longer-term selection against a broader population of potential mates, rather than only those intended for the next generation. Whether this helps balance short- and long-term response, and how that mate population should be defined, remain open questions.

## Conclusions

Multiple measures of genetic merit were theoretically derived over time. This study distinguishes additive genetic value, defined with a within-generation regression of individual’s genetic values on their allele dosages in a reference population, from breeding value, defined with a between-generation regression of expected progeny genetic values on parental allele dosages with clearly specified mate pools and parental roles. The two values coincide under the classical benchmark with a common reference and mate population, Hardy–Weinberg genotype frequencies, and no imprinting. They can differ with dominance and genetically differentiated mate pools, or with imprinting, even when female and male allele frequencies are equal. The Price equation connects the breeding-value expressions obtained by the between-generation regression to the change in mean genetic value under selection. Under this definition we obtain that breeding value equals twice GCA when both use the same mate population and mean progeny value.

These distinctions are relevant for animal and plant breeding programs with different reproductive groups, including purebred and crossbred populations. To properly interpret different genetic merit measures, the reference population, mate population, and parental role should be stated explicitly. Future work is to develop and test statistical procedures that account for these conditions when estimating the different measures of genetic merit.

## Acknowledgments

DLC acknowledges support from the Strategic Grant Plan of the Department of Science, University, and Knowledge Society 2022-2026 of the Government of Aragon, as well as from the Fundación Caja Inmaculada and the Aragón Agri-Food Institute (IA2) for funding the research stay during which this work was carried out. LV acknowledges support from Spanish Ministry of Science and Innovation grant PID2024-157480OB-I00. GG acknowledges support from the BBSRC ISP grants (BBS/E/RL/230001A, BBS/E/RL/230001B, and BBS/E/RL/230001C). The authors acknowledge use of generative AI tools for improving grammar and narrative flow. The authors carefully reviewed the final manuscript and take full responsibility for the accuracy and integrity of the content. For the purpose of open access, the authors have applied a CC BY public copyright license to any Author Accepted Manuscript version arising from this submission.

## Conflicts of interest

The authors state no conflicts of interest.

## Data availability

The Rubia Gallega dataset analyzed during the current study is proprietary to a private breeding program and is not publicly accessible. However, these data may be obtained from the corresponding author upon reasonable request. All other datasets used in this study were generated via simulation. The custom code used to generate the simulated data and perform all analyses is openly available on GitHub at: https://github.com/david20011999/Breeding-value-theory-with-genetically-differentiated-mate-pools-and-imprinting.git.

## Appendix

### A. Centering allele dosages, additive genetic values, and breeding values

The main text defines additive genetic value and breeding value using centered allele dosages. Here, we show that centering changes the regression intercept, but leaves the slope and fitted genetic values unchanged for a fixed reference population, equivalent with the widely used definition in Falconer (1981). This equivalence matters when estimating these quantities from data, because allele dosages can be centered in different ways (e.g., Strandén and Christensen 2011). We first consider the classical benchmark without imprinting, with Hardy–Weinberg genotype frequencies and *p*_*f*_ = *p*_*m*_ = *p*. For genotypes 0*/*0, 0*/*1 or 1*/*0, and 1*/*1 with genetic values *−a, d*, and *a*, the mean genetic value is *µ*_*G*_ = *a*(*p− q*) + 2*dpq* and the within-generation slope is *α*^*w*^ = *a* + *d*(*q −p*). With uncentered allele dosage *N*_1,*j*_, the regression intercept is *µ*_*G*_ *−*2*pα*^*w*^. Centering the dosage at its mean, 2*p*, gives intercept *µ*_*G*_ in (2). In either representation, additive genetic value is the fitted genetic value minus the mean genetic value in the reference population:

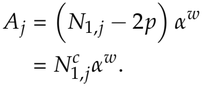

The same value can be written as a sum of allele effects. Using 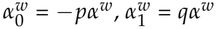, and *N*_0,*j*_ = 2 *− N*_1,*j*_ gives:

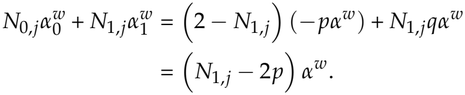

Thus, both expressions give the additive genetic values in Falconer (1981):

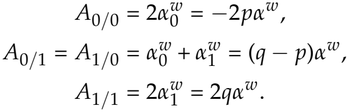

This invariance to centering holds for any fixed reference population. The Hardy–Weinberg assumption is needed here for the stated expression of *α*^*w*^. With imprinting, the same argument applies to the two origin-specific allele dosages in (6), each centered at its mean in the specified reference population.

For breeding value, centering likewise leaves the parent-to-offspring slope *b*_*s*_ and the between-generation coefficient 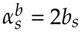 unchanged. For a parent *j* evaluated in role *s*, the one-locus model gives:

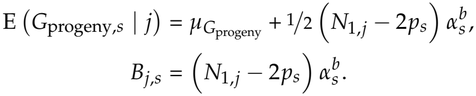

Here, *p*_*s*_ is the allele frequency in the evaluated parental pool, and 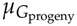 is the mean progeny genetic value across random pairings of the female and male pools. Using uncentered dosage changes the progeny-regression intercept to 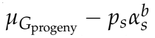, but leaves the breeding values unchanged. Under the classical benchmark without imprinting, *α*^*b*^ = *α*^*w*^ and *B*_*j*_ = *A*_*j*_. Changing the reference population or mate pool, however, can change these values; this is a different operation from centering a dosage within a fixed population.

### B. Derivation of the between-generation coefficient

We derive the between-generation coefficient *α*^*b*^ = 2*b* under undifferentiated and differentiated mate pools, using (5):

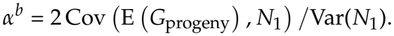

We use uncentered allele dosage because centering changes neither the covariance nor the variance (Appendix A). Pairing is random between pools and segregation is Mendelian. We first derive the coefficient without imprinting, then show how imprinting enters through expected progeny genetic value.

**Under undifferentiated mate pools**, the reference population and both mate pools have allele frequency *p* and Hardy– Weinberg genotype frequencies. Table B1 gives the expected progeny genetic value for each parental genotype *g*, averaged over the mate pool and Mendelian segregation. Averaging over parental genotypes gives the first two moments of allele dosage:

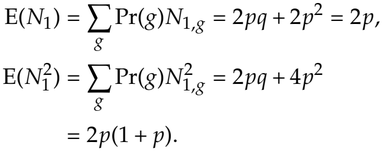

The mean progeny genetic value and the expected product of progeny expectation and parental allele dosage are:

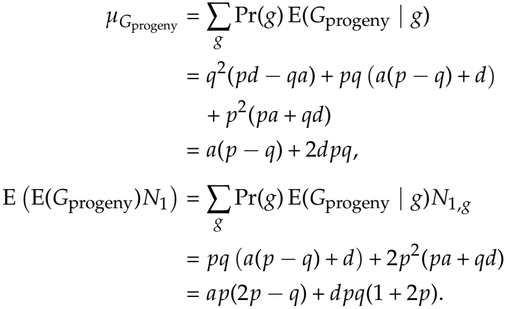

The covariance and variance therefore simplify to:

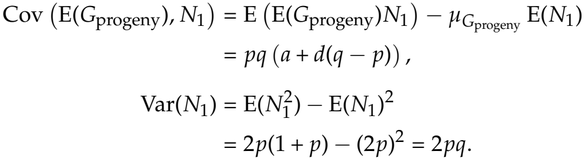

For 0 *< p* 1, the between-generation coefficient is:

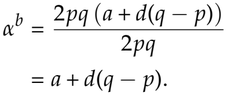

Thus, *α*^*b*^ = *α*^*w*^ under the classical benchmark without imprinting.

**With differentiated mate pools**, consider females evaluated as candidate parents with mates drawn randomly from the male pool. Table B2 separates the genotype frequencies of the evaluated females, which depend on *p*_*f*_, from their progeny expectations, which depend on the male mate frequency *p*_*m*_. The first two moments of female allele dosage are:

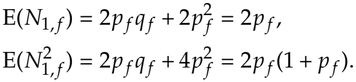

Averaging the progeny expectations over female genotypes gives:

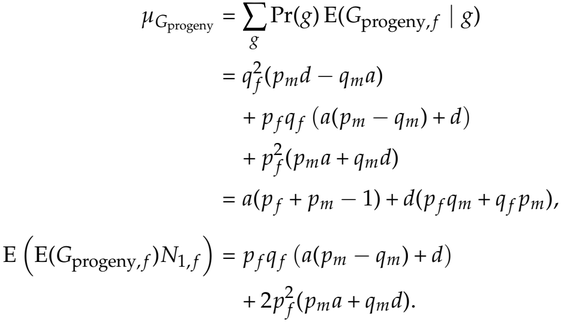

The covariance and variance are then:

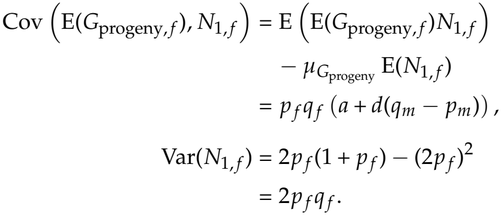

For 0 *< p*_*f*_ *<* 1, this gives:

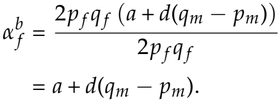

Exchanging the parental roles gives 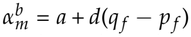. The co-efficient for each parental role therefore depends on the opposite-sex mate pool.

With imprinting, the progeny expectations for female parents with dosages 0, 1, and 2 become, respectively, *p*_*m*_(*d* + *i*) *−q*_*m*_*a*, 1*/*2*a*(*p*_*m*_ *−q*_*m*_) + 1*/*2*d* + 1*/*2*i*(*p*_*m*_ *−q*_*m*_), and *p*_*m*_*a* + *q*_*m*_(*d− i*). Their dependence on female allele dosage remains linear, with slope *b*_*f*_ = 1*/*2 (*a* + *d*(*q*_*m*_ *−p*_*m*_) *−i*). Exchanging the parental roles reverses the sign of the imprinting contribution, giving the coeffi-cients in (9) and (10):

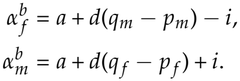

The tables use Hardy–Weinberg genotype frequencies to evaluate the moments. Because expected progeny genetic value is linear in parental allele dosage in this one-locus model, the same coefficients hold for other parental genotype frequencies with non-zero dosage variance. If that variance is zero, the regression slope cannot be estimated within the parental pool.

### C. Comparison of additive genetic values and breeding values for the two examples

We compare additive genetic values and breeding values for the two one-locus examples in Section Demonstration. Both examples use additive effect *a* = 1 and dominance effect *d* = 0.75, without imprinting in Figure 2 and with imprinting effect *i* = 0.5 in Figure 3. Genetic values and effects are expressed in arbitrary trait units. With undifferentiated mate pools, both sexes share a common reference population at *p*_*f*_ = *p*_*m*_ = *p* = 0.6. With differentiated mate pools, females and males have allele frequencies *p*_*f*_ = 0.3 and *p*_*m*_ = 0.9, and additive genetic values are calculated in their separate reference populations. All reference populations have Hardy–Weinberg genotype frequencies, and pairing is random between the female and male pools. Tables C1 and C2 give the corresponding values for each ordered genotype, written in maternal/paternal allele order.

In the first example, without imprinting, additive genetic values in the common reference population are 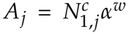, where 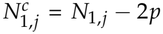 as visualised in Figure 2. With within-generation *α*^*w*^ = 0.85, the values for both females and males are: *A*_0*/*0_ = *−*1.02, *A*_0*/*1_ = *−*0.17, *A*_1*/*0_ = *−*0.17, and *A*_1*/*1_ = +0.68 (Table C1).

Under separate female and male reference populations, ad-ditive genetic values are 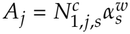, where 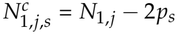 is centered within reference population *s*. With 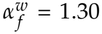 the values for females are: *A*_0*/*0_ = *−*0.78, *A*_0*/*1_ = +0.52, *A*_1*/*0_ = +0.52, and *A*_1*/*1_ = +1.82, while with 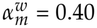 the values for males are: *A*_0*/*0_ = *−*0.72, *A*_0*/*1_ = *−*0.32, *A*_1*/*0_ = *−*0.32, and *A*_1*/*1_ = +0.08 (Table C1). The genetic value of each genotype is the same in females and males. The differences in additive genetic value arise from the different reference populations, which determine both the within-generation slopes and dosage centering. These values describe statistical decompositions within their respective references. Comparing individuals on a common reference scale requires the same reference population. The reference population must therefore be stated when reporting additive genetic value, just as the parental role and mate pool must be stated when reporting breeding value.

Under the classical benchmark, breeding values are 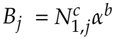 as visualised in Figure 2. The between-generation co-efficient *α*^*b*^ = 0.85 equals the within-generation slope *α*^*w*^, so breeding values equal additive genetic values (Table C1). The same numerical value therefore has two interpretations. As ad-ditive genetic value, it describes the individual itself: its fitted genetic value from the within-generation regression on allele dosage, minus the mean genetic value in the reference population. As breeding value, it describes the individual as a parent: twice the difference between the mean genetic value expected for its progeny and the mean progeny genetic value across all parents evaluated in the same role with that mate pool.

**Table B1.** Expected progeny genetic value at a single bi-allelic locus under the undifferentiated mate pools, without imprinting. The evaluated candidate parents and their mates have allele frequency *p* and Hardy–Weinberg genotype frequencies. Genotype frequencies refer to the evaluated candidate parents; expected progeny values average over random pairing and Mendelian segregation. Reciprocal heterozygotes are grouped because their progeny expectations are identical.

| Parental genotype *g* | *N*<sub>1,*g*</sub> | Frequency | E (*G*<sub>progeny</sub> \| *g*) | E (*G*<sub>progeny</sub> \| *g*) *N*<sub>1,*g*</sub> |
|---|---|---|---|---|
| 0/0 | 0 | *q*<sup>2</sup> | *pd* − *qa* | 0 |
| 0/1 & 1/0 | 1 | 2*pq* | <sup>1</sup>/2*a*(*p* − *q*) + <sup>1</sup>/2*d* | <sup>1</sup>/2*a*(*p* − *q*) + <sup>1</sup>/2*d* |
| 1/1 | 2 | *p*<sup>2</sup> | *pa* + *qd* | 2(*pa* + *qd*) |

**Table B2.** Expected progeny genetic value for female parents at a single bi-allelic locus, without imprinting, under differentiated mate pools. Evaluated females have allele frequency *p*_*f*_ and Hardy–Weinberg genotype frequencies; their mates are drawn randomly from the male pool with allele frequency *p*_*m*_. Genotype frequencies refer to the evaluated females; expected progeny values average over the male mate pool and Mendelian segregation. Reciprocal heterozygotes are grouped because their progeny expectations are identical.

| Female genotype *g* | *N*<sub>1,*g*,*f*</sub> | Frequency | E(*G*<sub>progeny,*f*</sub> \| *g*) | E(*G*<sub>progeny,*f*</sub> \| *g*)*N*<sub>1,*g*,*f*</sub> |
|---|---|---|---|---|
| 0/0 | 0 | *q*<sub>*f*</sub><sup>2</sup> | *p*<sub>*m*</sub>*d* − *q*<sub>*m*</sub>*a* | 0 |
| 0/1 & 1/0 | 1 | 2*p*<sub>*f*</sub>*q*<sub>*f*</sub> | 1/2*a*(*p*<sub>*m*</sub> − *q*<sub>*m*</sub>) + 1/2*d* | 1/2*a*(*p*<sub>*m*</sub> − *q*<sub>*m*</sub>) + 1/2*d* |
| 1/1 | 2 | *p*<sub>*f*</sub><sup>2</sup> | *p*<sub>*m*</sub>*a* + *q*<sub>*m*</sub>*d* | 2(*p*<sub>*m*</sub>*a* + *q*<sub>*m*</sub>*d*) |

**Table C1.**
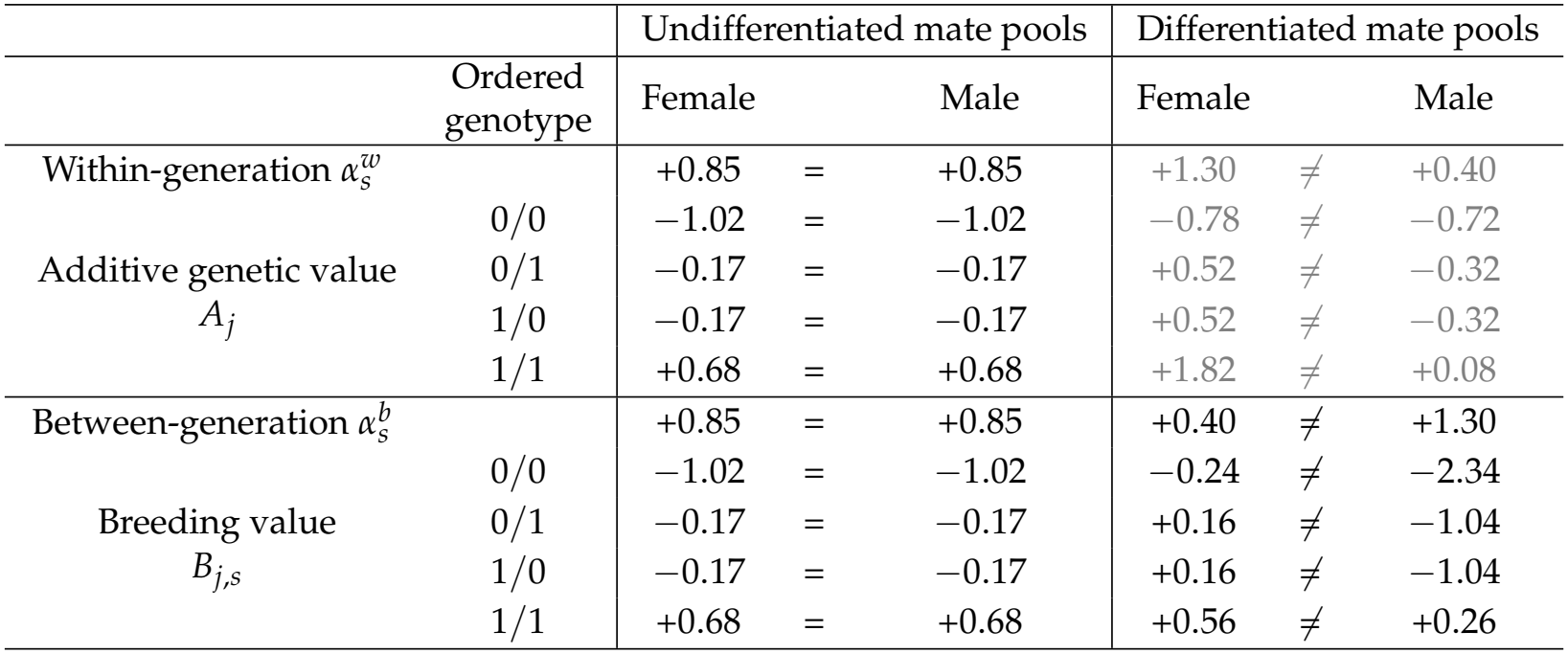
Comparison of additive genetic values and breeding values for each ordered genotype in Figure 2, without imprinting. Genotypes are written in maternal/paternal allele order. Reciprocal heterozygotes have identical values and are grouped in the figure. The trait is controlled by a single bi-allelic locus with additive effect *a* = 1, dominance effect *d* = 0.75, and no imprinting effect. Genetic values and effects are expressed in arbitrary trait units. Undifferentiated mate pools share the common reference frequency *p*_*f*_ = *p*_*m*_ = *p* = 0.6. Differentiated female and male pools have *p*_*f*_ = 0.3 and *p*_*m*_ = 0.9, with separate reference populations for additive genetic value. All reference populations have Hardy–Weinberg genotype frequencies. Additive genetic values are *A*_*j*_ = 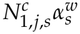; breeding values are 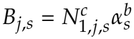 Dosage is centered at 2*p*_*s*_ in the indicated reference population or parental pool. Female and male columns identify reference populations for additive genetic value and parental roles for breeding value. Gray entries use separate female and male reference populations; their differences reflect the reference choice and do not compare individuals on a common reference scale.

|  |  | Undifferentiated mate pools |  |  | Differentiated mate pools |  |  |
|---|---|---|---|---|---|---|---|
|  | Ordered genotype | Female |  | Male | Female |  | Male |
| Within-generation α<sub>s</sub><sup>w</sup> |  | +0.85 | = | +0.85 | +1.30 | ≠ | +0.40 |
| Additive genetic value<br>A<sub>j</sub> | 0/0 | −1.02 | = | −1.02 | −0.78 | ≠ | −0.72 |
|  | 0/1 | −0.17 | = | −0.17 | +0.52 | ≠ | −0.32 |
|  | 1/0 | −0.17 | = | −0.17 | +0.52 | ≠ | −0.32 |
|  | 1/1 | +0.68 | = | +0.68 | +1.82 | ≠ | +0.08 |
| Between-generation α<sub>s</sub><sup>b</sup> |  | +0.85 | = | +0.85 | +0.40 | ≠ | +1.30 |
| Breeding value<br>B<sub>j,s</sub> | 0/0 | −1.02 | = | −1.02 | −0.24 | ≠ | −2.34 |
|  | 0/1 | −0.17 | = | −0.17 | +0.16 | ≠ | −1.04 |
|  | 1/0 | −0.17 | = | −0.17 | +0.16 | ≠ | −1.04 |
|  | 1/1 | +0.68 | = | +0.68 | +0.56 | ≠ | +0.26 |

**Table C2.**
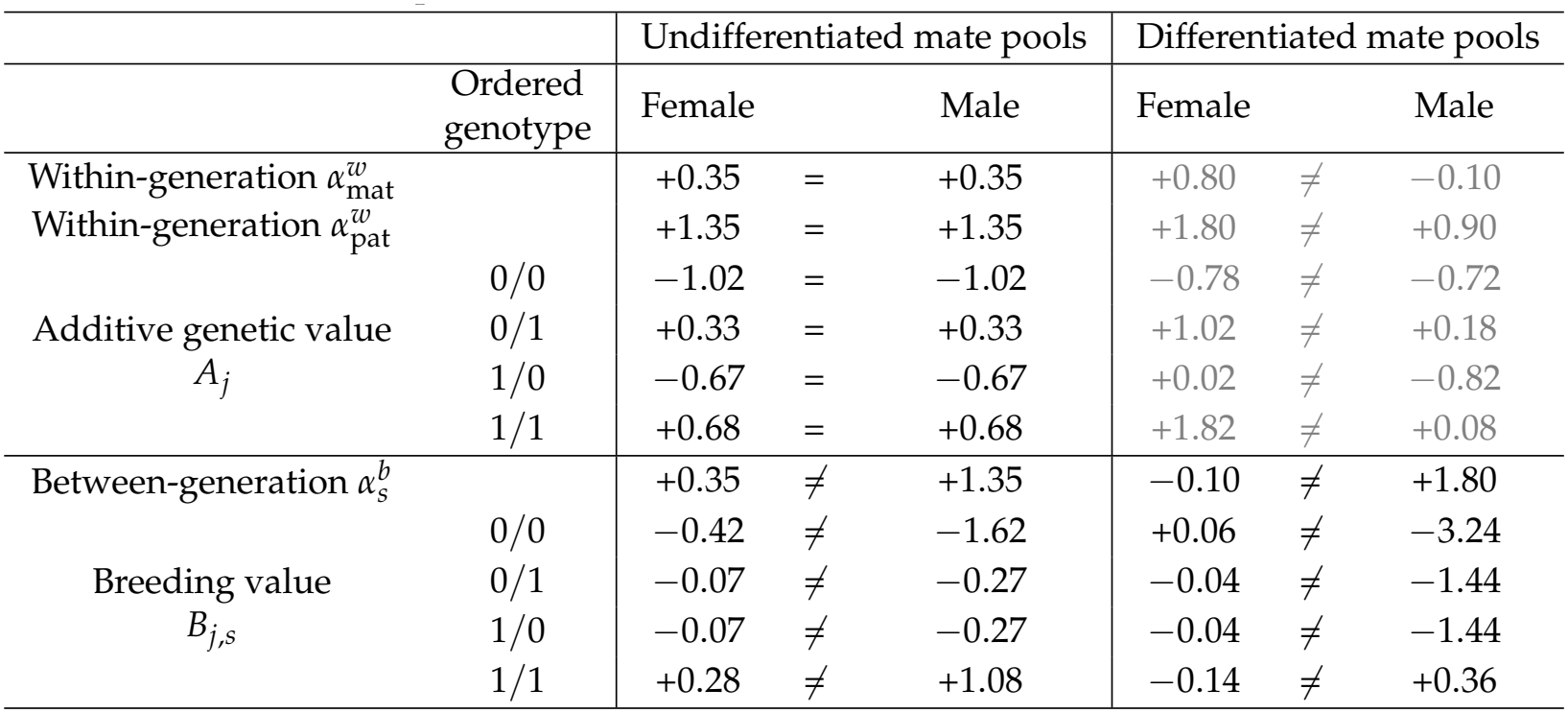
Comparison of additive genetic values and breeding values for each ordered genotype in Figure 3, with imprinting. Geno-types are written in maternal/paternal allele order. The trait is controlled by a single bi-allelic locus with additive effect *a* = 1, dominance effect *d* = 0.75, and imprinting effect *i* = 0.5. Genetic values and effects are expressed in arbitrary trait units. Undifferentiated mate pools share the common reference frequency *p*_*f*_ = *p*_*m*_ = *p* = 0.6. Differentiated female and male pools have *p*_*f*_ = 0.3 and *p*_*m*_ = 0.9, with separate reference populations for additive genetic value. All reference populations have Hardy–Weinberg genotype frequencies. Additive genetic values 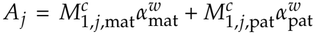. Both origin-specific allele dosages are centered at 0.6 in the common reference, at 0.3 in the female reference, and at 0.9 in the male reference. Breeding values are 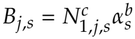, with total dosage centered at 2*p*_*s*_ in the evaluated parental pool. Female and male columns identify reference populations for additive genetic value and parental roles for breeding value. Gray entries use separate female and male reference populations; their differences reflect the reference choice and do not compare individuals on a common reference scale.

|  |  | Undifferentiated mate pools |  |  | Differentiated mate pools |  |  |
|---|---|---|---|---|---|---|---|
|  | Ordered genotype | Female |  | Male | Female |  | Male |
| Within-generation *α*<sup>*w*</sup><sub>mat</sub> |  | +0.35 | = | +0.35 | +0.80 | ≠ | −0.10 |
| Within-generation *α*<sup>*w*</sup><sub>pat</sub> |  | +1.35 | = | +1.35 | +1.80 | ≠ | +0.90 |
| Additive genetic value<br>*A*<sub>*j*</sub> | 0/0 | −1.02 | = | −1.02 | −0.78 | ≠ | −0.72 |
|  | 0/1 | +0.33 | = | +0.33 | +1.02 | ≠ | +0.18 |
|  | 1/0 | −0.67 | = | −0.67 | +0.02 | ≠ | −0.82 |
|  | 1/1 | +0.68 | = | +0.68 | +1.82 | ≠ | +0.08 |
| Between-generation *α*<sup>*b*</sup><sub>*s*</sub> |  | +0.35 | ≠ | +1.35 | −0.10 | ≠ | +1.80 |
| Breeding value<br>*B*<sub>*j*,*s*</sub> | 0/0 | −0.42 | ≠ | −1.62 | +0.06 | ≠ | −3.24 |
|  | 0/1 | −0.07 | ≠ | −0.27 | −0.04 | ≠ | −1.44 |
|  | 1/0 | −0.07 | ≠ | −0.27 | −0.04 | ≠ | −1.44 |
|  | 1/1 | +0.28 | ≠ | +1.08 | −0.14 | ≠ | +0.36 |

Under differentiated female and male mate pools, breeding values are 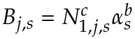 With 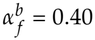 the values for females are: *B*_0*/*0_ = *−*0.24, *B*_0*/*1_ = +0.16, *B*_1*/*0_ = +0.16, and *B*_1*/*1_ =+0.56, while with 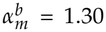 the values for males are: *B*_0*/*0_ = *−*2.34, *B*_0*/*1_ = *−*1.04, *B*_1*/*0_ = *−*1.04, and *B*_1*/*1_ = +0.26 (Table C1).

In this example, additive genetic values and breeding values rank genotypes in the same order within each sex because both coefficients are positive. Their magnitudes differ because the within-generation slope uses the specified reference population, whereas the between-generation coefficient uses the opposite-sex mate pool.

In the second example, with imprinting, additive genetic values are obtained using 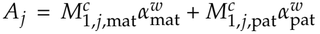. In the common reference population, both origin-specific allele dosages are centered at 0.6. With 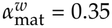 and 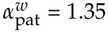, the values for females and males are: *A*_0*/*0_ = *−*1.02, *A*_0*/*1_ = +0.33,*A*_1*/*0_ = *−*0.67, and *A*_1*/*1_ = +0.68 (Table C2).

**Figure C1.**
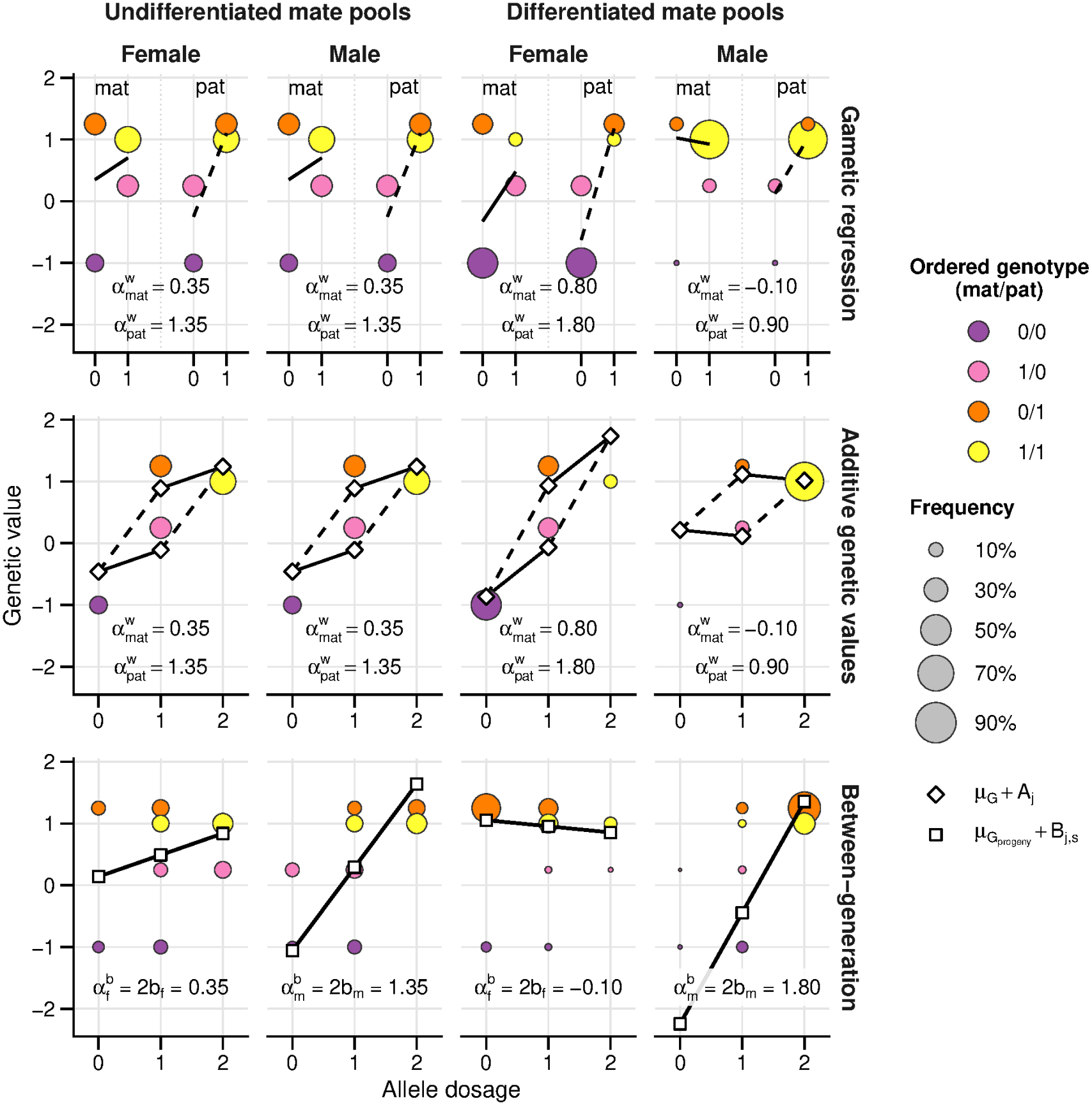
Within- and between-generation regressions in undifferentiated and differentiated mate pools, with imprinting. The trait is controlled by a single bi-allelic locus with additive effect *a* = 1, dominance effect *d* = 0.75, and imprinting effect *i* = 0.5. Undifferentiated mate pools have *p*_*f*_ = *p*_*m*_ = *p* = 0.6, while differentiated female and male pools have *p*_*f*_ = 0.3 and *p*_*m*_ = 0.9. Reference populations have Hardy–Weinberg genotype frequencies. Bubbles denote genetic values and colors identify ordered genotypes. In the top row, bubbles show individual genetic values *G*_*j*_ against origin-specific allele dosage *M*_1_ and recognizing the parental origin of alleles. Their area denotes genotype frequency in the female or male reference population. Solid and dashed lines show within-generation gamete regressions respectively giving 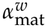 and 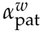. In the middle row, bubbles show individual genetic values *G*_*j*_ against allele dosage *N*_1_ and recognizing the parental origin of alleles. Their area denotes genotype frequency in the female or male reference population. Diamonds and segments show fitted genetic value *µ*_*G*_ + *A*_*j*_, with within-generation regressions giving additive genetic value *A*_*j*_. Solid segments change the maternally inherited allele while holding the paternally inherited allele fixed, and have slope 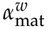 Dashed segments change the paternally inherited allele while holding the maternally inherited allele fixed, and have slope 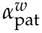. In the bottom row, bubbles show possible progeny genetic values against parental allele dosage *N*_1_. Their area denotes the frequency of each parent–progeny combination under random pairing. Squares and the solid line show fitted genetic value 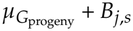, with slope 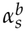, giving breeding value *B*_*j,s*_. See Table C2 for the actual additive genetic values and breeding values.

In the separate female reference population, both origin-specific allele dosages are centered at 0.3. With 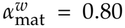 and 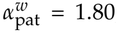, the values are: *A*_0*/*0_= *−*0.78, *A*_0_*/*_1_ = +1.02,*A*_1*/*0_ = +0.02, and *A*_1*/*1_ = +1.82. In the separate male reference population, both origin-specific allele dosages are centered at 0.9. With 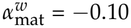 and 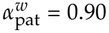, the values are: *A*_0*/*0_ = *−*0.72, *A*_0*/*1_ = +0.18, *A*_1*/*0_ = *−*0.82, and *A*_1*/*1_ = +0.08 (Table C2). As in the example without imprinting, these additive genetic values are defined within their respective reference populations. Each value includes both maternally and paternally inherited contributions. These inherited origins are distinct from the individual’s role as a female or male parent.

With imprinting, breeding values remain 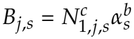, with dosage centered within the evaluated parental pool, but the between-generation coefficient depends on the parental role and the opposite-sex mate pool, as visualised in Figure 3 and Figure C1. With undifferentiated mate pools and 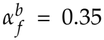, the values for female parents are: *B*_0*/*0_ = *−*0.42, *B*_0*/*1_ = *−*0.07,*B*_1*/*0_= *−*0.07, and *B*_1*/*1_ = +0.28, and with 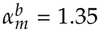 the values for male parents are: *B*_0*/*0_ = *−*1.62, *B*_0*/*1_ = *−*0.27, *B*_1*/*0_ = *−*0.27, and *B*_1*/*1_ = +1.08 (Table C2). In this example with imprinting, breeding values differ between female and male parents even with undifferentiated mate pools. The corresponding coefficients satisfy 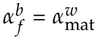 and 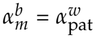, but this equality does not imply *A*_*j*_ =≠ *B*_*j,s*_. Additive genetic value weights the two inherited alleles by their origins, whereas breeding value weights total allele dosage by the individual’s parental role. Consequently, within each reference population *A*_0*/*1_ = *A*_1*/*0_, but within each parental role *B*_0*/*1_ = *B*_1*/*0_.

Under differentiated female and male mate pools, with 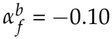 the values for females are: *B*_0*/*0_ = +0.06, *B*_0*/*1_ = *−*0.04, *B*_1*/*0_ = *−*0.04, and *B*_1*/*1_ = *−*0.14, while 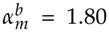 gives the values for males: *B*_0*/*0_ = *−* 3.24, *B*_0*/*1_ = *−* 1.44, *B*_1*/*0_ = *−* 1.44, and *B*_1*/*1_ = +0.36 (Table C2). In this example, the between-generation coefficient for female parents changes from +0.35 to *−*0.10 between undifferentiated and differentiated mate pools. This sign change reverses their genotype ranking by breeding value and the reciprocal heterozygotes remain tied.

### D. Response to selection and measurements of genetic merit

Under the classical benchmark of undifferentiated mate pools without imprinting, additive genetic value and breeding value coincide and provide the basis for predicting response with the breeder’s equation (Lush 1943). The Price equation provides a more general framework for decomposing evolutionary change of traits, into selection and transmission components (Price 1970, 1972). This framework connects animal breeding with evolutionary biology (Bijma 2020). Heywood (2005) used the Price equation to derive an exact generalization of the breeder’s equation and showed that the resulting partition of change in mean phenotype depends on whether genes, individuals, or mated pairs are treated as the parental units.

We use the Price equation to connect response to selection with the between-generation regression definition of breeding value within the one-locus model of the main text. This connection can be developed by calculating changes in expected progeny means from female and male allele frequencies, or by treating mating pairs as contributing units and evaluating their relative reproductive contributions and expected progeny genetic values. We show that the terms in the selection response that are linear in allele-frequency change are weighted by the between-generation coefficients, which can differ from the within-generation coefficients. This provides a selection-response justification for the between-generation definition of breeding value for a specified mate pool and parental role. For brevity and continuity with the main text, we develop only the simpler allele-frequency formulation under random pairing here and leave the pair-based derivation for a future publication.

Let *z*_*j*_ and 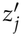 denote genetic values for a trait respectively in the parental and progeny generations. The change in mean genetic value between these generations is:

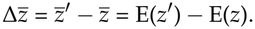

Price (1970, 1972) partitioned this change into selection and transmission components. For each parent *j*, let *w*_*j*_ denote its relative reproductive contribution (relative fitness, hereafter referred to as fitness), and let 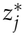 denote the expected genetic value of the progeny attributed to that parent. The transmission difference is 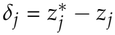 and relative fitness is normalized so that E(*w*) = 1. The Price equation then takes the form:

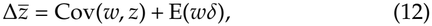

where the covariance and expectation are over the reference distribution of parents. The selection term Cov(*w, z*) measures the association between parental genetic value and fitness, while the transmission term E(*wδ*) accounts for differences between parental and expected progeny genetic values, weighted by fitness.

Using the normalization E(*w*) = 1, we can expand the transmission term as:

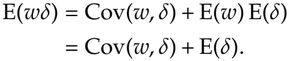

Substituting this expression into the Price equation, combining the covariance terms, and using 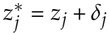 gives:

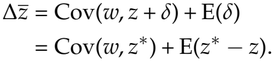

Rearranging isolates the covariance between fitness and expected progeny genetic value:

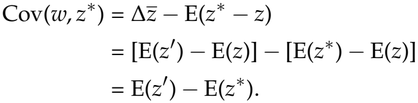

Thus, this covariance compares the mean genetic value in the progeny generation with the mean of the parent-specific progeny expectations over the reference parents. If selection changes only fitness and leaves these conditional progeny expectations unchanged, E(*z*^*\**^) is the mean progeny genetic value without (before) selection. Under this condition, E(*z*^*\**^ *−z*) represents the baseline change from parents to progeny in the absence of selection.

To evaluate the parental and expected progeny means in our one-locus model, let *p*_*f*_ and *p*_*m*_ denote the frequencies of allele 1 respectively in the female and male parental pools, with *q*_*f*_ = 1*− p*_*f*_ and *q*_*m*_ = 1 *−p*_*m*_. For the parental baseline, we assume

Hardy–Weinberg genotype frequencies and equal frequencies of reciprocal heterozygotes within each pool. The mean genetic values in the two parental pools are then:

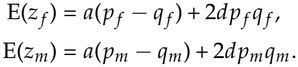

Giving equal weight to the female and male pools, we define the parental reference mean as:

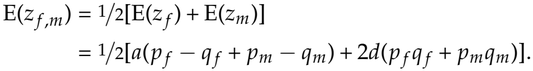

Under random pairing between these pools and Mendelian segregation, the expected progeny mean in the absence of selection is:

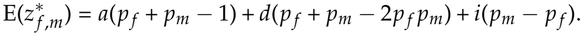

This progeny mean follows from the independent union of maternal and paternal gametes at frequencies *p*_*f*_ and *p*_*m*_.

After selection, let 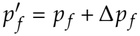 and 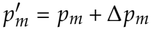 denote the frequencies of allele 1 respectively in the female and male parental pools. Here, Δ*p*_*f*_ and Δ*p*_*m*_ are the changes in allele frequency caused by selection within each pool. Under random pairing of the selected parents and Mendelian segregation, the expected progeny mean is obtained by substituting the allele frequencies under selection into the preceding expression:

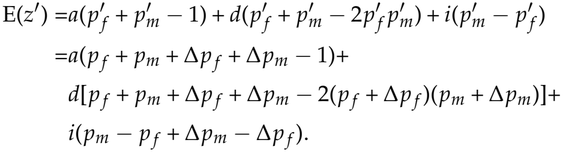

Substituting the progeny means before and after selection into the rearranged Price equation gives:

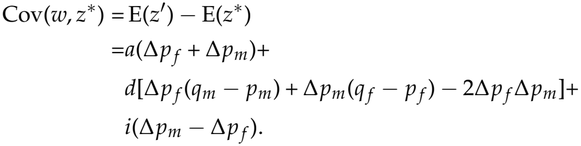

Collecting the terms in Δ*p*_*f*_ and Δ*p*_*m*_ gives:

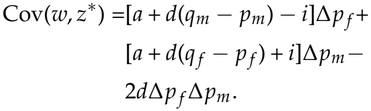

The coefficients of the two linear terms are 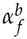 from (9) and 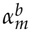 from (10), evaluated at the allele frequencies before selection. The interaction term *−*2*d*Δ*p*_*f*_ Δ*p*_*m*_ accounts for simultaneous changes in allele frequency in both pools.

To obtain the total change in mean genetic value under selection, we add the baseline change from parents to progeny in the absence of selection. Under the parental genotype-frequency assumptions stated above, this baseline is:

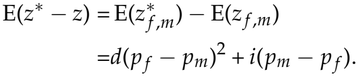

Substituting this baseline and the covariance expression into the Price equation gives:

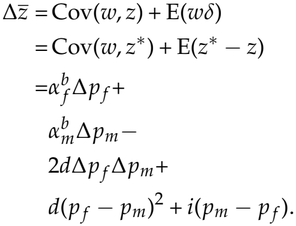

The terms involving Δ*p*_*f*_ and Δ*p*_*m*_ describe the change relative to the progeny mean without selection. The remaining terms describe the baseline change, which can be nonzero even when Δ*p*_*f*_ = Δ*p*_*m*_ = 0.

With the baseline change accounted for, the terms 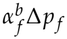 and 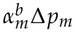 show how the between-generation coefficients weight the allele-frequency changes driven by selection. These are the same coefficients that define breeding value in the main text. For a specified mate pool, 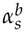 is twice the regression slope of expected progeny genetic value on parental allele dosage, as defined in (5). Multiplying this coefficient by the parent’s centered allele dosage gives 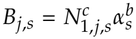. Because 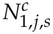 is centered on the allele fre-quency of the parent candidate pool, the mean 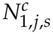 of selected candidates is 2Δ*p*_*s*_. Consequently, the average breeding value of selected females and males,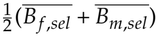, is the linear selec-tion component of the Price equation, 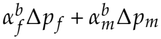. It also separates this change from the difference between parental and expected progeny means that can arise even without selection.

## Notes

https://github.com/david20011999/Breeding-value-theory-with-genetically-differentiated-mate-pools-and-imprinting.git

### Competing Interest Statement

The authors have declared no competing interest.

## Literature cited

Aguilar I, Misztal I, Johnson D, Legarra A, Tsuruta S, Lawlor T. 2010. Hot topic: a unified approach to utilize phenotypic, full pedigree, and genomic information for genetic evaluation of Holstein final score. Journal of dairy science. 93:743–752.

Álvarez-Castro JM. 2014. Dissecting genetic effects with imprinting. Frontiers in Ecology and Evolution. 2:51.

Alvarez-Castro JM, Carlborg O. 2007. A unified model for functional and statistical epistasis and its application in quantitative trait loci analysis. Genetics. 176:1151–1167.

Amer P, Crump R, Simm G. 1998. A terminal sire selection index for UK beef cattle. Animal Science. 67:445–454.

Antonios S, Legarra A, Astruc J, Rodríguez-Ramilo S, Vitezica Z. 2023. Partitioning of the genetic trends of French dairy sheep in Mendelian samplings and long-term contributions. Journal of Dairy Science. 106:6275–6287.

Ayres L, Bovenhuis H, Calus MP. 2025. A single-locus quantitative genetic model incorporating DNA methylation. Journal of Theoretical Biology. p. 112110.

Beef Improvement Federation. 1972. Guidelines for uniform beef improvement programs: Beef improvement federation recommendation. Technical Report 1020. United States Department of Agriculture, Extension Service. Washington, DC.

Bijma P. 2020. The Price equation as a bridge between animal breeding and evolutionary biology. Philosophical Transactions of the Royal Society B: Biological Sciences. 375:20190360.

Bonferroni C. 1936. Teoria statistica delle classi e calcolo delle probabilita. Pubblicazioni del R istituto superiore di scienze economiche e commericiali di firenze. 8:3–62.

Brabazon D, Callanan J, Nolan C. 2022. Imprinting of canine IGF2 and H19. Animal genetics. 53:108–118.

Calus MP, Schrooten C, Veerkamp RF. 2014. Genomic prediction of breeding values using previously estimated SNP variances. Genetics Selection Evolution. 46:52.

Casas E, Shackelford S, Keele J, Stone R, Kappes S, Koohmaraie M. 2000. Quantitative trait loci affecting growth and carcass composition of cattle segregating alternate forms of myostatin. Journal of animal science. 78:560–569.

Castillo-Bravo R, Fort A, Cashell R, Brychkova G, McKeown PC, Spillane C. 2022. Parent-of-origin effects on seed size modify heterosis responses in Arabidopsis thaliana. Frontiers in plant science. 13:835219.

Cheng Y, Rachagani S, Cánovas A, Mayes MS, Tait RG, Dekkers JC, Reecy JM. 2013. Body composition and gene expression QTL mapping in mice reveals imprinting and interaction effects. BMC genetics. 14:1–15.

Chenuil A, Crespin L, Pouyaud L, Berrebi P. 2004. Autosomal differences between males and females in hybrid zones: a first report from Barbus barbus and Barbus meridionalis (Cyprinidae). Heredity. 93:128–134.

Christensen OF, Legarra A, Lund MS, Su G. 2015. Genetic evaluation for three-way crossbreeding. Genetics Selection Evolution. 47:1–13.

Christensen OF, Madsen P, Nielsen B, Su G. 2014. Genomic evaluation of both purebred and crossbred performances. Genetics Selection Evolution. 46:23.

Cole J, VanRaden P. 2018. Symposium review: Possibilities in an age of genomics: The future of selection indices. Journal of dairy science. 101:3686–3701.

Davidson FA. 1925. Measuring the breeding value of dairy sires by the records of their first few advanced registry daughters. Bulletin 270. University of Illinois Agricultural Experiment Station. Urbana, Illinois.

Davis GP, Hetzel DJS, Corbet NJ, Scacheri S, Lowden S, Renaud J, Mayne C, Stevenson R, Moore SS, Byrne K. 1998. The mapping of quantitative trait loci for birth weight in a tropical beef herd. In: Piper L, editor, Proceedings of the 6th World Congress on Genetics Applied to Livestock Production. volume 26. pp. 441–444. Armidale, NSW, Australia. University of New England.

de Jong G, Powell O, Gorjanc G, Hickey JM, Gaynor RC. 2023. Comparison of genomic prediction models for general combining ability in early stages of hybrid breeding programs. Crop Science. 63:3338–3355.

De Koning DJ, Bovenhuis H, van Arendonk JA. 2002. On the detection of imprinted quantitative trait loci in experimental crosses of outbred species. Genetics. 161:931–938.

DeChiara TM, Robertson EJ, Efstratiadis A. 1991. Parental imprinting of the mouse insulin-like growth factor II gene. Cell. 64:849–859.

Falconer D. 1985. A note on Fisher’s ‘average effect’ and ‘average excess’. Genetics Research. 46:337–347.

Falconer DS. 1981. Introduction to quantitative genetics. Longman.

Ferguson-Smith AC, Cattanach B, Barton S, Beechey C, Surani M. 1991. Embryological and molecular investigations of parental imprinting on mouse chromosome 7. Nature. 351:667–670.

Fisher RA. 1919. The correlation between relatives on the supposition of Mendelian inheritance. Transactions of the Royal Society of Edinburgh. 52:399–433.

Fisher RA. 1922. On the interpretation of χ 2 from contingency tables, and the calculation of P. Journal of the royal statistical society. 85:87–94.

Fisher RA. 1930. The Genetical Theory of Natural Selection. Oxford: Clarendon Press.

Fisher RA. 1941. Average excess and average effect of a gene substitution. Annals of Eugenics. 11:53–63.

Fisher RA. 1958. The Genetical Theory of Natural Selection. 2nd edn. Oxford: Clarendon Press.

Flintham E. 2025. The evolution of sex-specific gene expression in polygenic traits. Journal of evolutionary biology. 38:939–951.

Flintham E, Lesaffre T, Otto S, Mullon C. 2026. Sexually antagonistic selection: a review of the theory and its implications. EcoEvoRxiv preprint.

Freeman A. 1984. Secondary traits: sire evaluation and the reproductive complex. Journal of Dairy Science. 67:449–458.

Gaynor RC, Gorjanc G, Hickey JM. 2021. AlphaSimR: an R package for breeding program simulations. G3. 11:jkaa017.

Giannoukakis N, Deal C, Paquette J, Goodyer CG, Polychronakos C. 1993. Parental genomic imprinting of the human IGF2 gene. Nature genetics. 4:98–101.

Graffelman J, Weir BS. 2018. On the testing of Hardy–Weinberg proportions and equality of allele frequencies in males and females at biallelic genetic markers. Genetic Epidemiology. 42:34–48.

Griffing B. 1956a. Concept of general and specific combining ability in relation to diallel crossing systems. Australian journal of biological sciences. 9:463–493.

Griffing B. 1956b. A generalised treatment of the use of diallel crosses in quantitative inheritance. Heredity. 10:31–50.

Grobet L, Royo Martin LJ, Poncelet D, Pirottin D, Brouwers B, Riquet J, Schoeberlein A, Dunner S, Ménissier F, Massabanda J et al. 1997. A deletion in the bovine myostatin gene causes the double–muscled phenotype in cattle. Nature genetics. 17:71–74.

Hager R, Cheverud JM, Leamy LJ, Wolf JB. 2008. Sex dependent imprinting effects on complex traits in mice. BMC evolutionary biology. 8:303.

Hayes BJ, Bowman PJ, Chamberlain AC, Verbyla K, Goddard ME. 2009. Accuracy of genomic breeding values in multi-breed dairy cattle populations. Genetics Selection Evolution. 41:1–9.

Henderson CR. 1975. Best linear unbiased estimation and prediction under a selection model. Biometrics. 31:423–447.

Henderson CR. 1984. Applications of Linear Models in Animal Breeding. University of Guelph. Guelph, Ontario, Canada.

Heywood JS. 2005. An exact form of the breeder’s equation for the evolution of a quantitative trait under natural selection. Evolution. 59:2287–2298.

Hill WG, Goddard ME, Visscher PM. 2008. Data and theory point to mainly additive genetic variance for complex traits. PLOS Genetics. 4:1–10.

Hill WG, Keightley PD. 1988. Interaction between molecular and quantitative genetics. In: Korver S, van der Steen HAM, van Arendonk JAM, Bakker H, Brascamp EW, Dommerholt J, editors, Advances in Animal Breeding: Proceedings of the World Symposium in Honour of Professor R. D. Politiek. pp. 41–55. Wageningen, The Netherlands. Pudoc.

Hivert V, Wray NR, Visscher PM. 2021. Gene action, genetic variation, and GWAS: A user-friendly web tool. PLOS Genetics. 17:1–9.

Hofmeister RJ, Cavinato T, Karimi R, van der Graaf A, Pajuste FD, Kronberg J, Taba N, Metspalu A, Esko T, Nelis M et al. 2025. Parent-of-origin effects on complex traits in up to 236,781 individuals. Nature. 646:647–656.

Hohenboken WD. 1985. The manipulation of variation in quantitative traits: a review of possible genetic strategies. Journal of Animal Science. 60:101–110.

Hu Y, Rosa GJ, Gianola D. 2015. A GWAS assessment of the contribution of genomic imprinting to the variation of body mass index in mice. BMC genomics. 16:1–15.

Ibáñez-Escriche N, Fernando RL, Toosi A, Dekkers JC. 2009. Genomic selection of purebreds for crossbred performance. Genetics Selection Evolution. 41:1–10.

Jaton C, Schenkel F, Malchiodi F, Sargolzaei M, Price C, Baes C, Miglior F. 2017. Genetic analysis for quality of frozen embryos produced by Holstein cattle donors in Canada. Journal of Dairy Science. 100:7320–7329.

Jiang J, Shen B, O’Connell JR, VanRaden PM, Cole JB, Ma L. 2017. Dissection of additive, dominance, and imprinting effects for production and reproduction traits in Holstein cattle. BMC genomics. 18:425.

Jirtle R. 1997. Geneimprint. http://www.geneimprint.com/site/genes-by-status.

Kambadur R, Sharma M, Smith TP, Bass JJ. 1997. Mutations in myostatin (GDF8) in double-muscled Belgian Blue and Piedmontese cattle. Genome research. 7:910–915.

Karaman E, Lund MS, Anche MT, Janss L, Su G. 2018. Genomic prediction using multi-trait weighted GBLUP accounting for heterogeneous variances and covariances across the genome. G3: Genes, Genomes, Genetics. 8:3549–3558.

Khaw HL, Ponzoni RW, Yee HY, bin Aziz MA, Mulder HA, Marjanovic J, Bijma P. 2016. Genetic variance for uniformity of harvest weight in Nile tilapia (Oreochromis niloticus). Aquaculture. 451:113–120.

Khramtsova EA, Davis LK, Stranger BE. 2019. The role of sex in the genomics of human complex traits. Nature Reviews Genetics. 20:173–190.

Köhler C, Wolff P, Spillane C. 2012. Epigenetic mechanisms underlying genomic imprinting in plants. Annual review of plant biology. 63:331–352.

Krätschmer I, Robinson MR. 2026. A quantitative genetic model for indirect genetic effects and genomic imprinting under random and assortative mating. Genetics. 232:iyag042.

Kropp J, Peñagaricano F, Salih S, Khatib H. 2014. Invited review: Genetic contributions underlying the development of preimplantation bovine embryos. Journal of dairy science. 97:1187–1201.

Lawson HA, Cheverud JM, Wolf JB. 2013. Genomic imprinting and parent-of-origin effects on complex traits. Nature Reviews Genetics. 14:609–617.

Lippincott WA. 1923. Poultry breeding records. Circular 99. Kansas State Agricultural College, Agricultural Experiment Station. Manhattan, Kansas.

Lopes MS, Bastiaansen JW, Harlizius B, Knol EF, Bovenhuis H. 2014. A genome-wide association study reveals dominance effects on number of teats in pigs. PloS one. 9:e105867.

López-Carbonell D, Legarra A, Altarriba J, Hervás-Rivero C, Sánchez-Díaz M, Varona L. 2024. Multiple trait bayesian analysis of partitioned genetic trends accounting for uncertainty in genetic parameters. an example with the Pirenaica and Rubia Gallega beef cattle breeds. Journal of Animal Breeding and Genetics.

Lucotte EA, Laurent R, Heyer E, Segurel L, Toupance B. 2016. Detection of allelic frequency differences between the sexes in humans: a signature of sexually antagonistic selection. Genome Biology and Evolution. 8:1489–1500.

Lucy M. 2001. Reproductive loss in high-producing dairy cattle: where will it end? Journal of dairy science. 84:1277–1293.

Lush JL. 1935. Progeny test and individual performance as indicators of an animal’s breeding value. Journal of Dairy Science. 18:1–19.

Lush JL. 1943. Animal breeding plans. Iowa State College Press. second edition.

Lynch M, Walsh B. 1998. Genetics and Analysis of Quantitative Traits. Sinauer Associates. Sunderland, Massachusetts, USA.

Mafra Fortuna G, Zumbach BJ, Johnsson M, Pocrnic I, Gorjanc G. 2024. Accounting for nuclear and mito genome in dairy cattle breeding - a simulation study. JDS Communications. 5:572–576.

Martinez-Castillero M, Then C, Altarriba J, Srihi H, López-Carbonell D, Díaz C, Martinez P, Hermida M, Varona L. 2021. Detection of genomic regions with pleiotropic effects for growth and carcass quality traits in the Rubia Gallega cattle breed. Animals. 11:1682.

McPherron AC, Lee SJ. 1997. Double muscling in cattle due to mutations in the myostatin gene. Proceedings of the National Academy of Sciences. 94:12457–12461.

Meuwissen TH, Hayes BJ, Goddard M. 2001. Prediction of total genetic value using genome-wide dense marker maps. genetics. 157:1819–1829.

Mrode R, Pocrnic I. 2023. Linear Models for the Prediction of the Genetic Merit of Animals. CABI. Wallingford, UK.

Neugebauer N, Luther H, Reinsch N. 2010a. Parent-of-origin effects cause genetic variation in pig performance traits. Animal. 4:672–681.

Neugebauer N, Räder I, Schild H, Zimmer D, Reinsch N. 2010b. Evidence for parent-of-origin effects on genetic variability of beef traits. Journal of animal science. 88:523–532.

Nishio M, Satoh M. 2015. Genomic best linear unbiased prediction method including imprinting effects for genomic evaluation. Genetics Selection Evolution. 47:1–10.

Norman H, Hutchison J, Wright J. 2008. Sire conception rate: New national AI bull fertility evaluation. AIPL Res. Rep. SCR1 (7-08).

O’Brien EK, Wolf JB. 2019. Evolutionary quantitative genetics of genomic imprinting. Genetics. 211:75–88.

O’Doherty AM, MacHugh DE, Spillane C, Magee DA. 2015. Genomic imprinting effects on complex traits in domesticated animal species. Frontiers in Genetics. 6:156.

Pacheco HA, Rezende FM, Peñagaricano F. 2020. Gene mapping and genomic prediction of bull fertility using sex chromosome markers. Journal of Dairy Science. 103:3304–3311.

Pardo AM, Casanova D, Rubio N, Andere C, Rodríguez E, Corva PM. 2023. An insight into population structure and genetic progress of Argentinean Holstein cattle. Journal of Animal Breeding and Genetics. 140:376–389.

Price GR. 1970. Selection and covariance. Nature. 227:520–521.

Price GR. 1972. Extension of covariance selection mathematics. Annals of Human Genetics. 35:485–490.

Purcell S, Neale B, Todd-Brown K, Thomas L, Ferreira MA, Bender D, Maller J, Sklar P, De Bakker PI, Daly MJ et al. 2007. PLINK: a tool set for whole-genome association and population-based linkage analyses. The American journal of human genetics. 81:559–575.

Reik W, Dean W, Walter J. 2001. Epigenetic reprogramming in mammalian development. Science. 293:1089–1093.

Roughsedge T, Amer P, Thompson R, Simm G. 2005a. Development of a maternal breeding goal and tools to select for this goal in UK beef production. Animal Science. 81:221–232.

Roughsedge T, Amer P, Thompson R, Simm G. 2005b. Genetic parameters for a maternal breeding goal in beef production. Journal of Animal Science. 83:2319–2329.

Sae-Lim P, Komen H, Kause A, Van Arendonk J, Barfoot A, Martin K, Parsons J. 2012. Defining desired genetic gains for rainbow trout breeding objective using analytic hierarchy process. Journal of animal science. 90:1766–1776.

Sanchez MP, Escouflaire C, Baur A, Bottin F, Hozé C, Boussaha M, Fritz S, Capitan A, Boichard D. 2023. X-linked genes influence various complex traits in dairy cattle. BMC genomics. 24:338.

Schaeffer L, Kennedy B, Gibson J. 1989. The inverse of the gametic relationship matrix. Journal of Dairy Science. 72:1266–1272.

Scott RJ, Spielman M, Bailey J, Dickinson HG. 1998. Parent-of-origin effects on seed development in Arabidopsis thaliana. Development. 125:3329–3341.

Shi R, Chen Z, Su G, Luo H, Liu L, Guo G, Wang Y. 2023. Genomic prediction of service sire effect on female reproductive performance in Holstein cattle: A comparison between different methods, validation population and marker densities. Journal of Animal Breeding and Genetics. 140:355–365.

Spencer HG. 2002. The correlation between relatives on the sup-position of genomic imprinting. Genetics. 161:411–417.

Sprague GF, Tatum LA. 1942. General vs. specific combining ability in single crosses of corn. Agronomy Journal. 34:923–932.

Strandén I, Christensen OF. 2011. Allele coding in genomic evaluation. Genetics Selection Evolution. 43:25.

Su G, Guldbrandtsen B, Aamand GP, Strandén I, Lund MS. 2014. Genomic relationships based on X chromosome markers and accuracy of genomic predictions with and without X chromosome markers. Genetics Selection Evolution. 46:1–12.

Sun C, VanRaden PM, Cole JB, O’Connell JR. 2014. Improvement of prediction ability for genomic selection of dairy cattle by including dominance effects. PloS one. 9:e103934.

Suzuki S, Renfree MB, Pask AJ, Shaw G, Kobayashi S, Kohda T, Kaneko-Ishino T, Ishino F. 2005. Genomic imprinting of IGF2, p57KIP2 and PEG1/MEST in a marsupial, the tammar wallaby. Mechanisms of development. 122:213–222.

Swanson-Wagner RA, DeCook R, Jia Y, Bancroft T, Ji T, Zhao X, Nettleton D, Schnable PS. 2009. Paternal dominance of trans-eQTL influences gene expression patterns in maize hybrids. science. 326:1118–1120.

Tier B, Sölkner J. 1993. Analysing gametic variation with an animal model. Theoretical and applied genetics. 85:868–872.

Tiezzi F, Maltecca C. 2015. Accounting for trait architecture in genomic predictions of US Holstein cattle using a weighted realized relationship matrix. Genetics Selection Evolution. 47:24.

Toro MA, Varona L. 2010. A note on mate allocation for dominance handling in genomic selection. Genetics Selection Evolution. 42:1–9.

Triantaphyllopoulos KA, Ikonomopoulos I, Bannister AJ. 2016. Epigenetics and inheritance of phenotype variation in live-stock. Epigenetics & chromatin. 9:1–18.

Tucci V, Isles AR, Kelsey G, Ferguson-Smith AC, Bartolomei MS, Benvenisty N, Bourc’his D, Charalambous M, Dulac C, Feil R et al. 2019. Genomic imprinting and physiological processes in mammals. Cell. 176:952–965.

Twomey AJ, Cromie AR, McHugh N, Berry DP. 2020. Validation of a beef cattle maternal breeding objective based on a cross-sectional analysis of a large national cattle database. Journal of Animal Science. 98:skaa322.

Uffelmann E, Huang QQ, Munung NS, De Vries J, Okada Y, Martin AR, Martin HC, Lappalainen T, Posthuma D. 2021. Genome-wide association studies. Nature Reviews Methods Primers. 1:59.

Van Laere AS, Nguyen M, Braunschweig M, Nezer C, Collette C, Moreau L, Archibald AL, Haley CS, Buys N, Tally M et al. 2003. A regulatory mutation in IGF2 causes a major QTL effect on muscle growth in the pig. Nature. 425:832–836.

VanRaden PM. 2008. Efficient methods to compute genomic predictions. Journal of dairy science. 91:4414–4423.

Varona L, Legarra A, Toro MA, Vitezica ZG. 2018. Non-additive effects in genomic selection. Frontiers in genetics. 9:341678.

Vitezica ZG, Legarra A, Toro MA, Varona L. 2017. Orthogonal estimates of variances for additive, dominance, and epistatic effects in populations. Genetics. 206:1297–1307.

Vitezica ZG, Varona L, Elsen JM, Misztal I, Herring W, Legarra A. 2016. Genomic BLUP including additive and dominant variation in purebreds and F1 crossbreds, with an application in pigs. Genetics Selection Evolution. 48:1–8.

Walsh B, Lynch M. 2018. Evolution and Selection of Quantitative Traits. Oxford University Press. Oxford, UK.

Wang H, Misztal I, Aguilar I, Legarra A, Muir W. 2012. Genome-wide association mapping including phenotypes from relatives without genotypes. Genetics research. 94:73–83.

Wei M, van der Werf JH. 1994. Maximizing genetic response in crossbreds using both purebred and crossbred information. Animal Science. 59:401–413.

Werner CR, Gaynor RC, Sargent DJ, Lillo A, Gorjanc G, Hickey JM. 2023. Genomic selection strategies for clonally propagated crops. Theoretical and Applied Genetics. 136.

Wientjes YC, Bijma P, Calus MP, Zwaan BJ, Vitezica ZG, van den Heuvel J. 2022. The long-term effects of genomic selection: 1. response to selection, additive genetic variance, and genetic architecture. Genetics Selection Evolution. 54:19.

Wolf JB, Hager R, Cheverud JM. 2008. Genomic imprinting effects on complex traits: a phenotype-based perspective. Epigenetics. 3:295–299.

Wright S. 1921. Systems of mating. I. the biometric relations between parent and offspring. Genetics. 6:111–123.

Yapp WW. 1925. Transmitting ability of dairy sires. Journal of Animal Science. pp. 90–92.

Yoshida GM, Yáñez JM, de Oliveira CAL, Ribeiro RP, Lhorente JP, de Queiroz SA, Carvalheiro R. 2018. Mate selection allows changing the genetic variability of the progeny while optimizing genetic response and controlling inbreeding. Aquaculture. 495:409–414.

Zhang Z, Liu J, Ding X, Bijma P, de Koning DJ, Zhang Q. 2010. Best linear unbiased prediction of genomic breeding values using a trait-specific marker-derived relationship matrix. PloS one. 5:e12648.

